# Enhanced functionality of low-affinity CD19 CAR T-cells is associated with activation priming and a polyfunctional cytokine phenotype

**DOI:** 10.1101/2020.09.22.291831

**Authors:** Ilaria M. Michelozzi, Eduardo Gomez Castaneda, Ruben V.C. Pohle, Ferran Cardoso Rodriguez, Jahangir Sufi, Pau Puigdevall Costa, Meera Subramaniyam, Efstratios Kirtsios, Ayad Eddaoudi, Si Wei Wu, Aleks Guvenel, Jonathan Fisher, Sara Ghorashian, Martin A. Pule, Christopher J. Tape, Sergi Castellano, Persis J. Amrolia, Alice Giustacchini

## Abstract

We recently described a low-affinity second-generation CD19 chimeric antigen receptor (CAR) CAT that showed enhanced expansion, cytotoxicity, and anti-tumour efficacy compared to the high-affinity (FMC63 based) CAR used in Tisagenlecleucel, in pre-clinical models. Furthermore, CAT demonstrated an excellent toxicity profile, enhanced *in vivo* expansion, and long-term persistence in a Phase I clinical study. To understand the molecular mechanisms behind these properties of CAT CAR T-cells, we performed a systematic *in vitro* characterization of the transcriptomic (RNA-seq) and protein (CyTOF) changes occurring in T-cells expressing low-affinity *vs* high-affinity CD19 CARs following stimulation with CD19-expressing cells. Our results show that CAT CAR T-cells exhibit enhanced activation to CD19 stimulation and a distinct transcriptomic and protein profile, with increased activation and cytokine polyfunctionality compared to FMC63 CAR T-cells. We demonstrate that the enhanced functionality of low-affinity CAT CAR T-cells is a consequence of an antigen-dependent priming induced by residual CD19-expressing B-cells present in the manufacture.

## Introduction

T-cells genetically engineered to express CD19 chimeric antigen receptors (CAR T-cells) have shown remarkable efficacy in relapsed/refractory (r/r) B-cell malignancies leading to clinical licensing for r/r B-cell acute lymphoblastic leukaemia (B-ALL) and Non-Hodgkin Lymphoma^1^. Despite this success, several safety and efficacy hurdles remain^2^. CAR T-cells can trigger potent immune responses leading to transient but potentially life-threatening inflammatory events, such as cytokine release syndrome (CRS) and neurotoxicity^3,4^. CAR T-cell engraftment and expansion is a prerequisite for efficacy. Not all CD19 CAR T-cell products persist *in vivo* and long-term immune surveillance appears critical for preventing disease relapse, particularly in ALL. Thus, the design of versatile CAR T-cells, capable of balancing safety and efficacy, is contingent on our understanding of the molecular mechanisms underlying CAR T-cell function. The engagement of CARs to their cognate antigens results in the activation of CAR T-cells and promotes their rapid expansion as well as their differentiation into distinct T-cell subsets, mediating tumour cytolysis (effector cells) and providing long-lasting protection (memory cells). As tumour cell recognition by CAR T-cells relies on the binding of the CAR’s single-chain variable fragment (scFv) to its epitope, fine-tuning the affinity of CARs to their antigens has become a strategy to modulate the strength of CAR T-cell responses^5^.

The affinity of CARs is determined by their binding kinetics and the rates at which they associate to and dissociate from their targets. The optimal affinity of a CAR is likely to vary depending on a number of factors, including the CAR design, the CAR expression levels and the antigen density on the target cells^6,7^. Chimeric immunoreceptors have an activation ceiling above which increasing the binding affinity does not improve T-cell activation but can rather result in T-cell exhaustion^8^. In contrast, by reducing CAR scFv affinity, the strength of the T-cell signal can be modulated, so that CAR T-cells discriminate different levels of antigen expression. CARs exhibiting slower antigen-association rates to ErbB2, EGFR and CD123 targets showed reduced activation in response to low antigen concentrations, favouring differential targeting of tumour cells overexpressing the target *vs* normal tissue expressing the same target at physiological levels^8–10^. We have recently described a novel low-affinity CD19 CAR (CAT), with epitope, structure and stability similar to the widely used FMC63, but characterized by faster rates of antigen dissociation, leading to an overall 40-fold reduction of its affinity^11^. Pre-clinical testing of CAT CAR T-cells has revealed greater antigen-specific cytotoxicity, higher proliferation both *in vitro* and *in vivo* and more potent *in vivo* anti-tumour activity when compared to FMC63. In a Phase I clinical trial in patients with high-risk treatment-refractory paediatric B-ALL, CAT CAR T-cells resulted in lower toxicity in terms of severe CRS and displayed greater expansion than that reported for the FMC63 based tisagenlecleucel as well as excellent persistence^11^. These results were recently confirmed in a multicenter Phase I trial in r/r adult B-ALL^12^.

In this regard, while CAT CAR T-cells have been designed to explore the possibility to prevent excessive activation and reduce CRS and neurotoxicity risks, enhancements in their cytotoxicity, expansion and persistence were unexpected^7^ and the underlying molecular mechanisms to these functional phenotypes remain unknown.

The molecular mechanisms through which the fine-tuning of CARs affinity influences CAR T-cell phenotypes and functions are largely unknown. The interaction between T-cell receptor (TCR) and the peptide-MHC complex offers however some insight on how immunoreceptors’ affinity can dramatically influence T-cell functions^13^. Faster target off-rates in TCRs allow a single peptide-MHC complex to serially trigger several TCRs, resulting in amplified and sustained T-cell activation^13^. Similar to TCRs, low-affinity CARs may lead to enhanced T-cell activation and decreased exhaustion.

Herein we perform a systematic *in vitro* characterization of the molecular and biochemical changes occurring in CAR T-cells, comparing a low-affinity CD19 CAR (CAT) with a high-affinity one (FMC63). By combining bulk RNA sequencing with single-cell mass cytometry analyses (cytometry by time of flight, CyTOF), we show that the expression of the CAT CAR induces an antigen-dependent priming in response to low concentrations of CD19-expressing B-cells found in the manufacture product. Upon antigen stimulation, we identified distinct molecular features downstream of CAT CAR activation responsible for enhancing CAT CAR T-cell functional response.

## Results

### Generation and quality assessments of low- (CAT) and high- (FMC63) affinity CD19 CARs from healthy donors

To dissect the molecular mechanisms behind the functional differences observed between low- (CAT) and high- (FMC63) affinity CD19 CAR T-cells^11^, we interrogated the transcriptional and protein expression profiles of T-cells lentivirally (LV) transduced with CARs differing only in their scFv (Supplementary Fig. 1a). We performed bulk transcriptomic analyses (RNA-seq) to identify CAR T-cell distinct gene expression signatures and mass cytometry analyses (CyTOF) to model differences in their downstream signalling at a single-cell resolution^14^. RNA-seq and CyTOF readouts from untransduced (UNTR) controls and T-cells LV transduced to express CAT or FMC63 CD19 CARs from healthy donors (HD1-HD27, Supplementary Table 1) were compared at baseline and following stimulation with CD19+ ALL cell line NALM6 (unstimulated and stimulated conditions, respectively), as schematized in Fig. 1a. Importantly, we ruled out significant differences in the transduction of the two CARs by fluorescence activated cell sorting (FACS), assessing the percentage of mCherry+ T-cells (LV fluorescent reporter) across donors and experimental conditions. These ranged between 11.50 - 93.80% (median 39.1%) in FMC63 and 13.40 - 93.60% (median 36.25%) in CAT (Fig. 1b, left) and were thus comparable among individual HDs. In agreement, we found similar transgene expression levels between the 2 CARs by measuring mCherry mean fluorescent intensity (MFI), a proxy for CARs expression (Fig. 1b, right) and by quantifying CARs surface expression levels (Fig. 1c) and number of integrated vector copies (VCN) (Supplementary Fig. 1b). Finally, we assessed CAT and FMC63 CAR T-cell products for their CD4:CD8 ratios and memory T-cell subsets composition, as these characteristics can both affect CAR T-cells persistence and anti-tumour activity^15,16^. While no difference in CD4:CD8 ratios was observed in the absence of antigen stimulation (Fig. 1d), the proportion of memory subsets differed in unstimulated FMC63 and CAT CAR T-cells, as measured by FACS at 10 days post-transduction. CAT CAR T-cells exhibited a significant increase in the fraction of central memory T-cells (T_CM_, CD62+CD45RA-) as compared to both UNTR and FMC63 conditions (Fig. 1e, left). The increase in T_CM_ was largely at the expense of T effector memory cells (T_EM_, CD62-CD45RA-) and effector memory re-expressing CD45RA (T_EMRA_, CD62-CD45RA+), whose proportion was significantly reduced in CAT *vs* FMC63 CAR T-cells and in both CARs as compared to UNTR control (Fig. 1e, middle and right). These assessments confirmed that our CAR T-cell products were suitable to investigate the molecular features of CAT and FMC63 CAR T-cells.

**Fig. 1:**
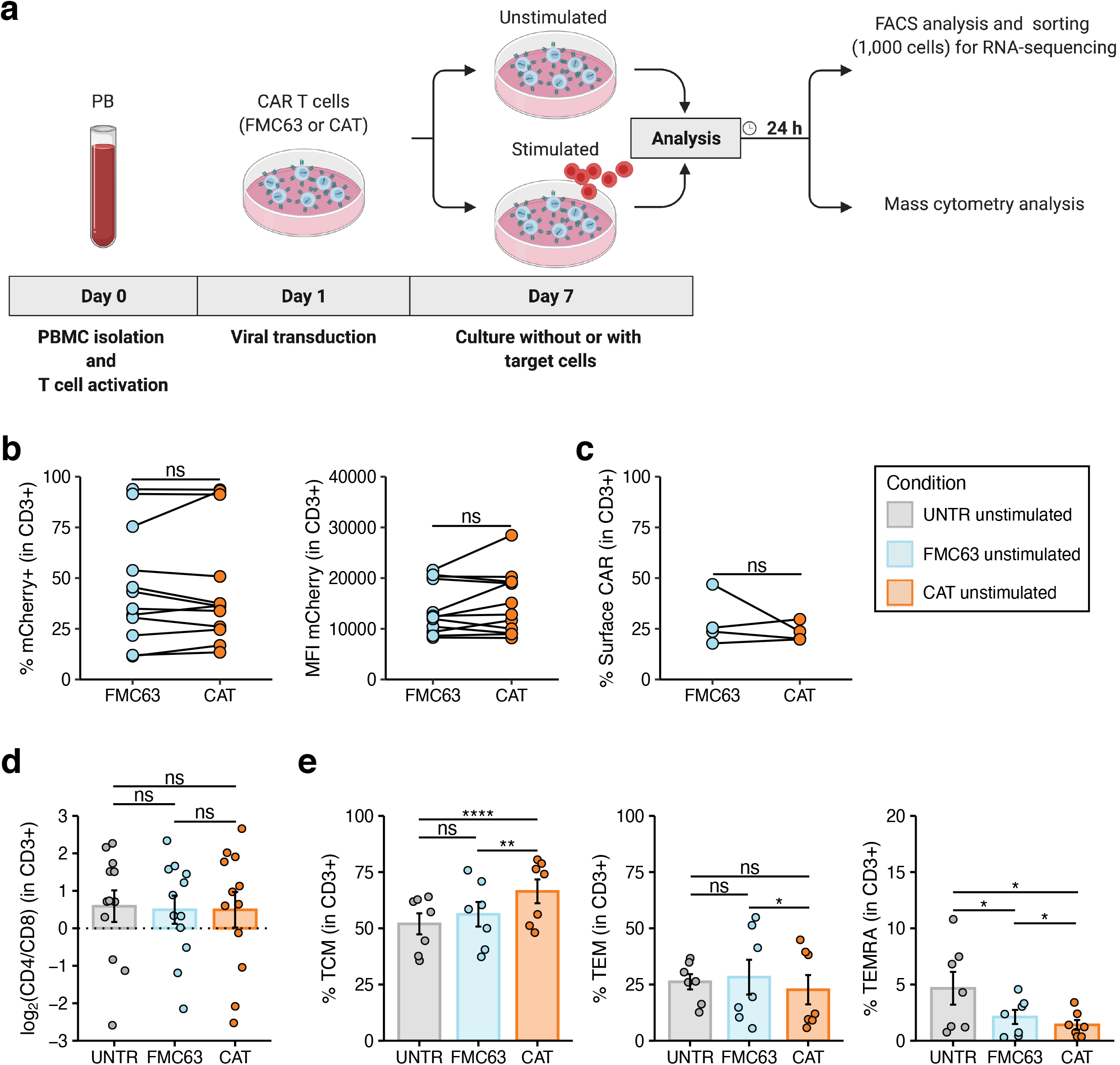
Generation and phenotypic characterisation of CAR T-cells from HD-PBMCs. **a** Experimental workflow. Peripheral Blood Mononuclear Cells (PBMCs) were isolated from HDs and LV transduced to express CD19 CAR construct (FMC63 or CAT) following overnight activation with CD3/CD28 beads. Six days after transduction, CAR T-cells were cultured without (unstimulated) or with target cells (NALM6) at 1:1 ratio (stimulated). Unstimulated and stimulated cells were analysed by flow cytometry and sorted for RNA-sequencing 24 hours post-stimulation. Mass cytometry analysis was performed on unstimulated and stimulated cells at 24 h post-stimulation. Activated UNTR T-cells were used as a control throughout the experiment. **b** (left) Spaghetti plots showing transduction levels of CAR T-cells as percentage of mCherry+ (in CD3+) and (right) as MFI of mCherry in unstimulated transduced T-cells measured by FACS 7 days post-transduction. Lines connect results from individual donors (n= 12 HDs, n = 3 independent experiments). **c** Spaghetti plot showing the percentage of surface CAR expression (in CD3+) in unstimulated transduced T-cells measured by FACS 10 days post-transduction. Lines connect results from individual donors (n = 4 HDs, n = 1 independent experiment). **d** Variation (log2 fold change) of CD4 and CD8 proportion in unstimulated UNTR T-cells and FMC63 and CAT CAR T-cells measured by FACS 7 days post-transduction. The dotted horizontal line (0) represents the conditions in which CD4=CD8. Data represent mean ± se (n = 12 HDs, n = 3 independent experiments). **e** (left) Barplots showing the percentage of TCM (CD45RA-CD62L+), (middle) TEM (CD45RA-CD62L-) and (right) TEMRA (CD45RA+CD62L-) in unstimulated CD3+ UNTR T-cells and FMC63 and CAT CAR T-cells measured by FACS 10 days post-transduction. Data represent mean ± se (n = 7 HDs, n = 2 independent experiments). **b-e** Statistical significance was calculated by Paired t-test. *P < 0.05, **P < 0.01, ****P < 0.0001. Each experimental condition is indicated by a specific colour code (UNTR light grey, FMC63 light blue, CAT orange).

### Low-affinity CD19 CAT CARs display higher activation priming during CAR T cell manufacture

We next performed bulk RNA-seq of FACS-sorted UNTR T-cells (CD3+) and FMC63 or CAT CAR T-cells (CD3+mCherry+) expanded for 4 days with or without stimulation with NALM6 at 24 h post-stimulation. Transcriptomics confirmed that mCherry was an accurate proxy for CAR transgene expression levels, as evidenced by the significant positive correlation between the MFI of mCherry by FACS and the normalized RNA-seq counts aligning to the scFv region of each of the 2 CARs (Supplementary Fig. 2a). Similarly, the proportion of CD4 and CD8 T-cells detected by FACS was in line with the *CD4* and *CD8* mRNA levels (Supplementary Fig. 2b, c). Principal component analysis (PCA) on the 500 most variable expressed genes (top 100 genes shown in Supplementary Fig. 2d), distributed samples according to a T-cell activation gradient (PC1, from UNTR to CAR activated samples) (Fig. 2a). The majority of variance in gene expression across experimental conditions was explained by CD19-mediated CAR activation. As expected, UNTR T-cells not expressing any CARs were largely unaffected by antigen stimulation (Fig. 2a). Similarly, PCA on protein expression from CyTOF, based on earth mover’s distance (EMD) scores (a sensitive measure of multivariate changes in protein levels)^17^, in the same experimental conditions and timepoints than the gene expression, followed a similar gradient of sample activation, shifting from UNTR samples to antigen stimulated CAR T-cells on PC1 (Fig. 2b and Supplementary Fig. 3a). The most variable genes upregulated upon stimulation with NALM6 in both FMC63 and CAT CAR T-cells included genes involved in T-cell activation (*IL2RA, GZMB*) and proliferation (*PCNA, LDHA*), which are expressed at very low levels in control T-cells (Supplementary Fig. 2d). Consistent with this, the highest protein expression variation was from markers of T-cell activation (CD25, NFAT1, HLA-DR) and proliferation (pRB) (Supplementary Fig. 3a). Of note, both RNA and protein analyses unstimulated CAR T-cells had an intermediate RNA/protein expression profile between UNTR and stimulated CAR T-cells. This suggests that CAR expression on its own, in absence of antigen stimulation, induces basal T-cell activation (Fig. 2a-b). Importantly, unstimulated CAT samples appear closer to antigen stimulated CAR T-cells than unstimulated FMC63, as well as having less overlap with UNTR samples suggesting unstimulated CAT CAR T-cells show a degree of pre-activation even prior to antigen stimulation.

**Fig. 2:**
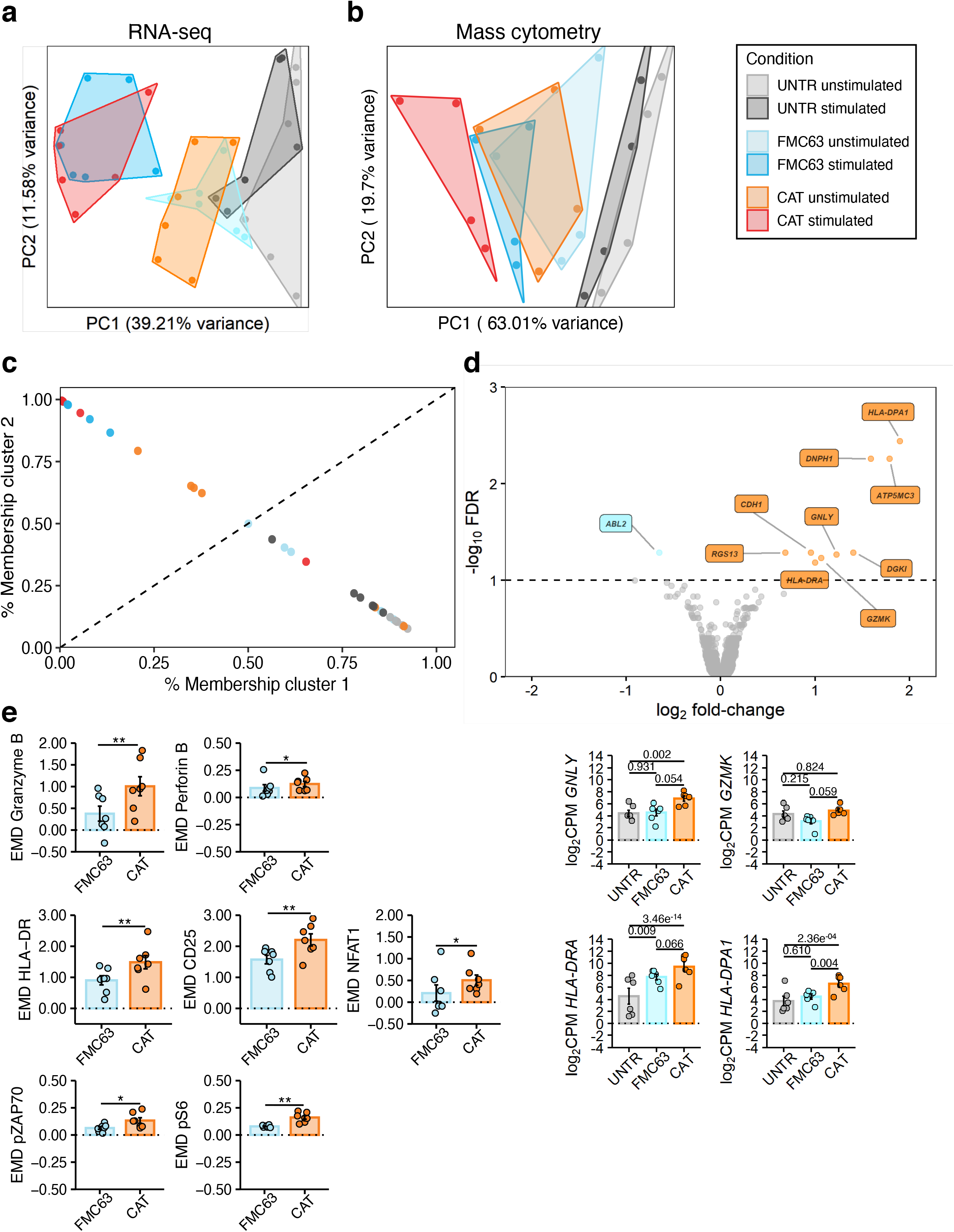
RNA-seq and mass cytometry analyses of unstimulated untransduced and CAR-transduced T-cells. **a** PCA of the top 500 variable genes from RNA-seq analysis across all the experimental conditions (n = 6 HDs, n = 2 independent experiments). **b** PCA of mass cytometry EMD scores computed at 24 h post-stimulation in CD3+ cells across all the experimental conditions (n = 4 HDs, n = 1 independent experiment). **c** Fuzzy clustering analysis of RNA-seq data across all experimental conditions (n = 6 HDs, n = 2 independent experiments). **d**(top) Volcano plot showing DE genes between unstimulated FMC63 and CAT CAR T-cells. The dashed horizontal line represents the statistical significance threshold (FDR < 0.1). (bottom) The barplots show the expression of selected DE genes (FDR < 0.1) in unstimulated untransduced and transduced T-cells. Data represent mean ± se (n = 6 HDs, n = 2 independent experiments). **e** The barplots show the expression of mass cytometry EMD scores for Granzyme B, Perforin B, HLA-DR, CD25, NFAT1, pZAP70 and pS6 in unstimulated CAR T-cells at 24 h upon stimulation. The data shown are normalized to stimulated CD3+ UNTR T-cells. The dotted horizontal line (0) represents the expression of a specific marker in unstimulated CD3+ UNTR T-cells. Data represent mean ± se (n = 7 HDs, n = 2 independent experiments). Statistical significance was calculated by Paired t-test. *P < 0.05, **P < 0.01. **a-e** Each experimental condition is indicated by a specific colour code (Unstimulated conditions = UNTR light grey, FMC63 light blue, CAT orange; stimulated conditions = UNTR grey, FMC63 blue, CAT red).

To gain further insights into the intermediate activation state observed in unstimulated CAT, we fuzzy clustered genes by their gene expression and classified individual samples into more than one cluster to resolve intermediate cell states and trajectories (Fig. 2c)^18^. We identified two clusters, highly enriched for either inactive T-cells (cluster 1, which includes UNTR samples) or antigen-activated CAR T-cells (cluster 2, which includes stimulated CAR T samples) (Fig. 2c). Notably, the probability of unstimulated CAT samples of belonging to cluster 2 (activated CAR T-cells) was substantially higher than that of unstimulated FMC63 (4/6 HDs in CAT *vs* 0/6 HDs inFMC63), further evidencing the functional proximity between unstimulated CAT and activated CAR T-cells. In this regard, using gene set enrichment analyses (GSEA) on the Hallmark collection, we confirmed that while antigen stimulated CAT and FMC63 CAR T-cells have similar enrichment for most of the gene sets involved in immune functions and cell proliferation, in the absence of antigen stimulation CAT CAR T-cells are uniquely enriched for T-cell activation pathways (Myc Targets V1, Hedghog signalling, Inflammatory response, Interferon α response) (Supplementary Fig. 3b).

To investigate whether the introduction of high- or low-affinity CD19 CARs can differentially affect gene expression patterns in T-cells, we directly compared the transcriptome of FMC63 and CAT CAR T-cells, in absence of antigen stimulation. Following differential gene expression (DGE) analysis, we found that only 10 genes were significantly DE between these two conditions (FDR < 0.1, Supplementary Table 2), 9 of which were upregulated in CAT *vs* FMC63 CAR T-cells. Among those, we found genes involved in cytotoxicity (*GNLY, GZMK*) and markers of T-cell activation such as MHC class II molecules (MHCII) (*HLA-DRA* and *HLA-DPA*) (Fig. 2d). This supports our previous observation of gene expression in unstimulated CAT resembling more that of antigen activated CAR T-cells than gene expression in unstimulated FMC63 (Fig. 2c). Mass cytometry analyses using a panel of antibodies against markers of T-cell activation (Supplementary Table 3), confirmed that while unstimulated FMC63 and CAT CAR T-cells have similar EMD scores for many of the proteins investigated (Supplementary Table 5a), unstimulated CAT CAR T-cells have significantly stronger activation priming with higher expression of markers of T-cell activation (HLA-DR, CD25 and NFAT1), pro-inflammatory (Granzyme B, Perforin B) and stimulatory/activation-related (GM-CSF, IL-17A) cytokines and increased phosphorylation of the TCR/CAR CD3ζ chain (pZAP70) and MTOR downstream effector (pS6) (Fig.2e and Supplementary Fig. 3c).

Altogether, these results show that unstimulated CAT CAR T-cells, prior to antigenic stimulation, have more pronounced T-cell activation priming than FMC63 CAR T-cells.

### CD19 stimulation of low-affinity CAT CAR T-cells results in a distinct transcriptomic and protein profile with increased activation/proliferation over high-affinity FMC63 CAR T-cells

While antigen-independent CAR activation, also known as tonic signalling, has been historically associated with CAR T-cell accelerated differentiation and exhaustion^19–21^, recent data show that the induction of CAR T-cell priming, by either 4-1BB-based tonic signaling^22^ or by low-dose hypomethylating agents^23^, can lead to CAR T enhanced anti-tumour functions *in vivo*. In light of these observations, we next wanted to assess the molecular impact of the “activation priming” observed in CAT CAR T-cells on their molecular response to antigen stimulation. As the superior cytotoxicity of CAT CAR T-cells over FMC63 CAR T-cells has been previously shown in functional assays^11^, we focused on characterizing their distinct molecular profiles upon exposure to antigenic stimulation. Upon stimulation, we found a slight but statistically significant reduction of CD4:CD8 ratio in CAT as compared to FMC63 (Fig. 3a). This variation was largely attributable to a relative decrease of the CD8+ fraction (and increase of CD4+) in FMC63 CAR T-cells after antigen stimulation (Supplementary Fig. 4a). Interestingly, when looking at the memory T-cell subsets composition after antigenic stimulation, we found that CAT CAR T-cells continued to exhibit a higher proportion of T_CM_ as compared to FMC63 (Fig. 3b), while no differences were observed in the expression of exhaustion markers (PD1, TIM3 and LAG3) between the two CAR constructs (Fig. 3c).

**Fig. 3:**
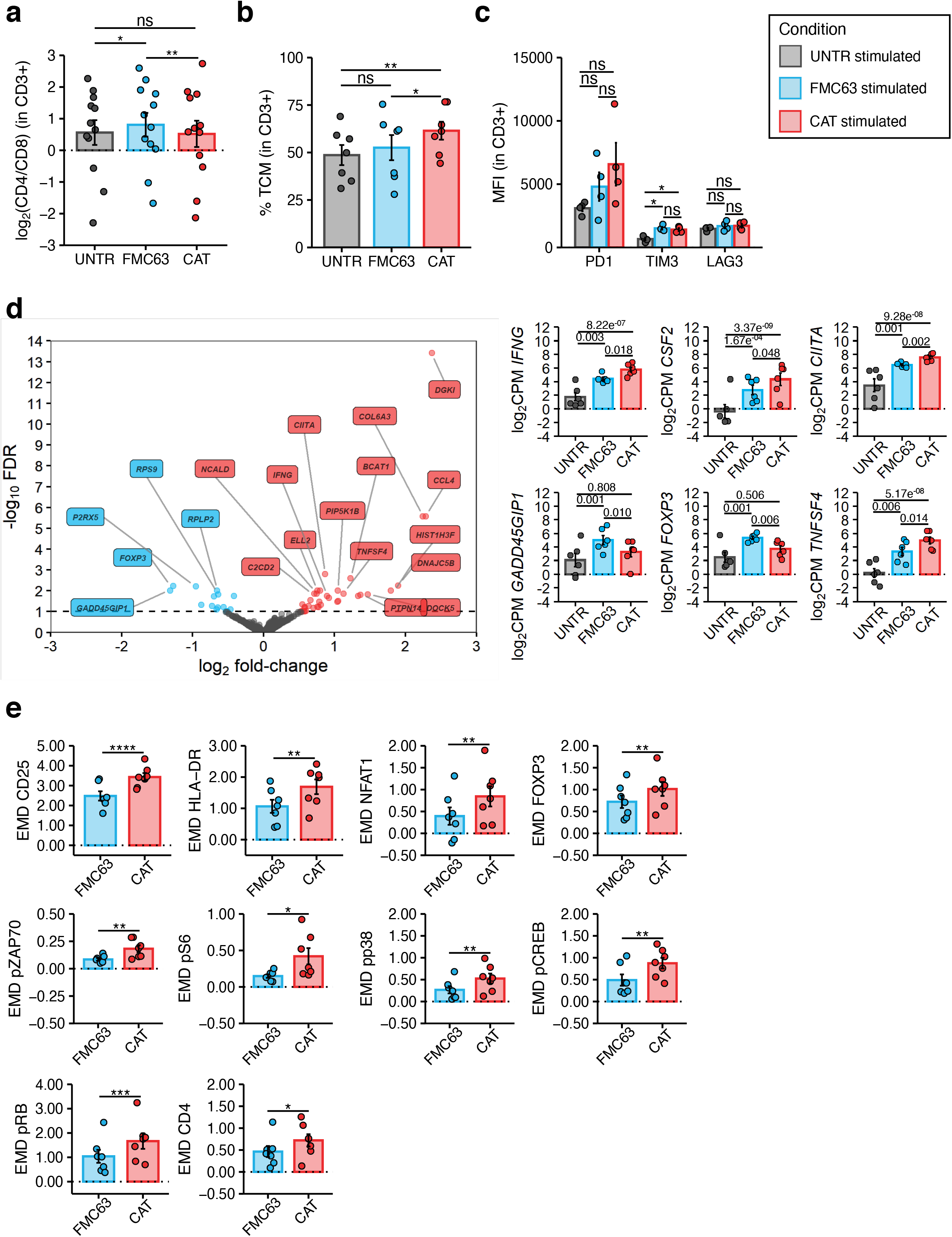
Phenotypic and molecular characterisation of stimulated CAR-transduced T-cells. **a** Variation (log2 fold change) of CD4 and CD8 proportion in stimulated UNTR T-cells and FMC63 and CAT CAR T-cells measured by FACS (n = 12 HDs, n = 3 independent experiments). The dotted horizontal line (0) represents the conditions in which CD4=CD8. **b** Barplot showing the percentage of TCM (CD45RA-CD62L+) in stimulated CD3+ UNTR T-cells and FMC63 and CAT CAR T-cells measured by FACS 96h post-antigen stimulation (n = 7 HDs, n = 2 independent experiments). **c** Barplots showing the expression of T-cell exhaustion markers (PD1, TIM3 and LAG3) as MFI in stimulated CD3+ UNTR T-cells and FMC63 and CAT CAR T-cells measured by FACS 96h post-antigen stimulation (n = 4 HDs, n = 1 independent experiment). **d** (left) Volcano plot showing DE genes between FMC63 and CAT CAR T-cells upon NALM6 co-culture. The dashed horizontal line represents the statistical significance threshold (FDR < 0.1). (right) The barplots show the expression of selected DE genes (FDR < 0.1) in stimulated UNTR T-cells and in CAR T-cells (n = 6 HDs, n = 2 independent experiments). **e** The barplots show the expression of mass cytometry EMD scores for CD25, HLA-DR, NFAT1, FOXP3, pZAP70, pS6, pp38, pCREB, pRB and CD4 in stimulated CAR T-cells at 24 h upon stimulation. The data shown are normalized to stimulated CD3+ UNTR T-cells. The dotted horizontal line (0) represents the expression of a specific marker in stimulated CD3+ UNTR T-cells. (n = 7 HDs, n = 2 independent experiments). **a-e** Barplots show mean ± se. Each experimental condition is indicated by a specific colour code (UNTR grey, FMC63 blue, CAT red). **a, b, c, e** Statistical significance was calculated by Paired t-test. *P < 0.05, **P < 0.01, ***P < 0.001, ****P < 0.0001.

Importantly, the stronger basal activation observed in CAT CAR T-cells did not prevent an even stronger molecular response when exposed to CD19-expressing NALM6, as shown by the increase in the expression of proliferation and cytotoxic/stimulatory markers, relatively to the unstimulated constructs (Supplementary Fig. 5 and 6). Consistent with this, after stimulation with NALM6, we identified 51 DE genes, 35 of which were upregulated in CAT compared to FMC63 (Fig. 3d and Supplementary Table 4). CD19 stimulation in CAT CAR T-cells also led to significantly augmented expression of immune stimulatory/proliferation cytokines (*IFNG, CSF2, CXCL8*), and IFN-γ responsive genes (*CIITA*) (Fig. 3d and Supplementary Fig. 4b). Conversely, CAT CAR T-cells displayed significantly decreased expression of the genes encoding for CRIF1 (*GADD45GIP1*), an inhibitor of cell cycle progression, and for FOXP3, the Treg-associated transcription factor known to be only transiently expressed in the initial stages of Th1 response and rapidly downregulated afterwards^24,25^ (Fig. 3d). These results suggest that following antigenic stimulation CAT CAR T-cells show a different transcriptomic profile resulting in stronger activation and proliferation than stimulated FMC63. In addition, CAT CAR T-cells showed increased expression of the *TNFSF4* gene (Fig. 3d), encoding for the ligand of the T-cell co-stimulatory receptor OX40 (OX40L). While OX40L is mainly expressed by antigen presenting cells to promote T-cell activation during antigenic stimulation, it is also expressed in activating T-cells, where it leads to a homotypic OX40L-OX40 signalling axis promoting T-cell longevity and memory differentiation^26^. Further, we also noted the increased expression of the chemoattractants *CCL4* and *CCL3L1* and genes involved in cell migration (*FLT1, DOCK5*) and focal adhesion (*COL6A3*) in stimulated CAT CAR T-cells (Supplementary Fig. 4b). Importantly, the secretion of CCL4 and CCL3 by activated T-cells has been shown to mediate T-cell attraction toward one another, promoting T-cell clusters formation and synaptic cytokines exchange coordinating immune responses^27–30^. Single cell transcriptomic studies have recently revealed that subsets of CAR T with elevated expression of CCL3 and CCL4 are associated to longer persistence *in vivo^31^* and achievement of complete remission^32^.

These observations were further substantiated at the protein level. Indeed, CAT CAR T-cells exhibited a marked increase in the expression (as measured by EMD score) of markers of T cell activation (CD25) (Fig. 3e). Moreover, the augmented gene expression of the MHCII trans-activator CIITA observed in CAT, resulted in a corresponding increase of HLA-DR protein (Fig. 3e)^33,34^. When measuring the CAR T-cell intracellular signalling, CAT CAR T-cells showed enhanced phosphorylation of the effectors of the TCR/CAR CD3ζ chain (pZAP70, pp38) and increased expression of their downstream transcription factors (NFAT1, pCREB, FOXP3). CAT CAR T-cells also exhibited significant upregulation of the Target of Rapamycin Complex 1 (mTORC1) downstream effectors (pS6 and pRB), both involved in cell proliferation and protein translation, in line with the previously reported CAT CAR T cells increased proliferative capacity over FMC63^11^.

Our analysis shows that the increased “activation priming” observed in CAT CAR T-cells over FMC63 resulted in an even stronger T-cell activation gene expression and signaling profile when CAT CAR T-cells were exposed to antigenic stimulation. These observations are in line with the CAT CAR T cells enhanced cytotoxic functional properties previously reported^11^.

### The enhanced functionality of low-affinity CD19 CAR T-cells is associated with cytokine polyfunctionality upon antigen stimulation

We next assessed CAR T-cell functional phenotypes by measuring intracellular cytokines levels in individual cells by mass cytometry. The overall protein intensities measured by EMD scores indicated an increased expression of effector cytokines (Granzyme B, IFN-γ and TNF-α) and of immune stimulatory molecules (GM-CSF and IL-2) in CAT CAR T as compared to FMC63 (Figure 4a). No upregulation was observed for Th2/immune-modulatory cytokines (IL-4, IL-5 and TGF-ß) and for Perforin B (Supplementary Table 5b).

**Fig. 4:**
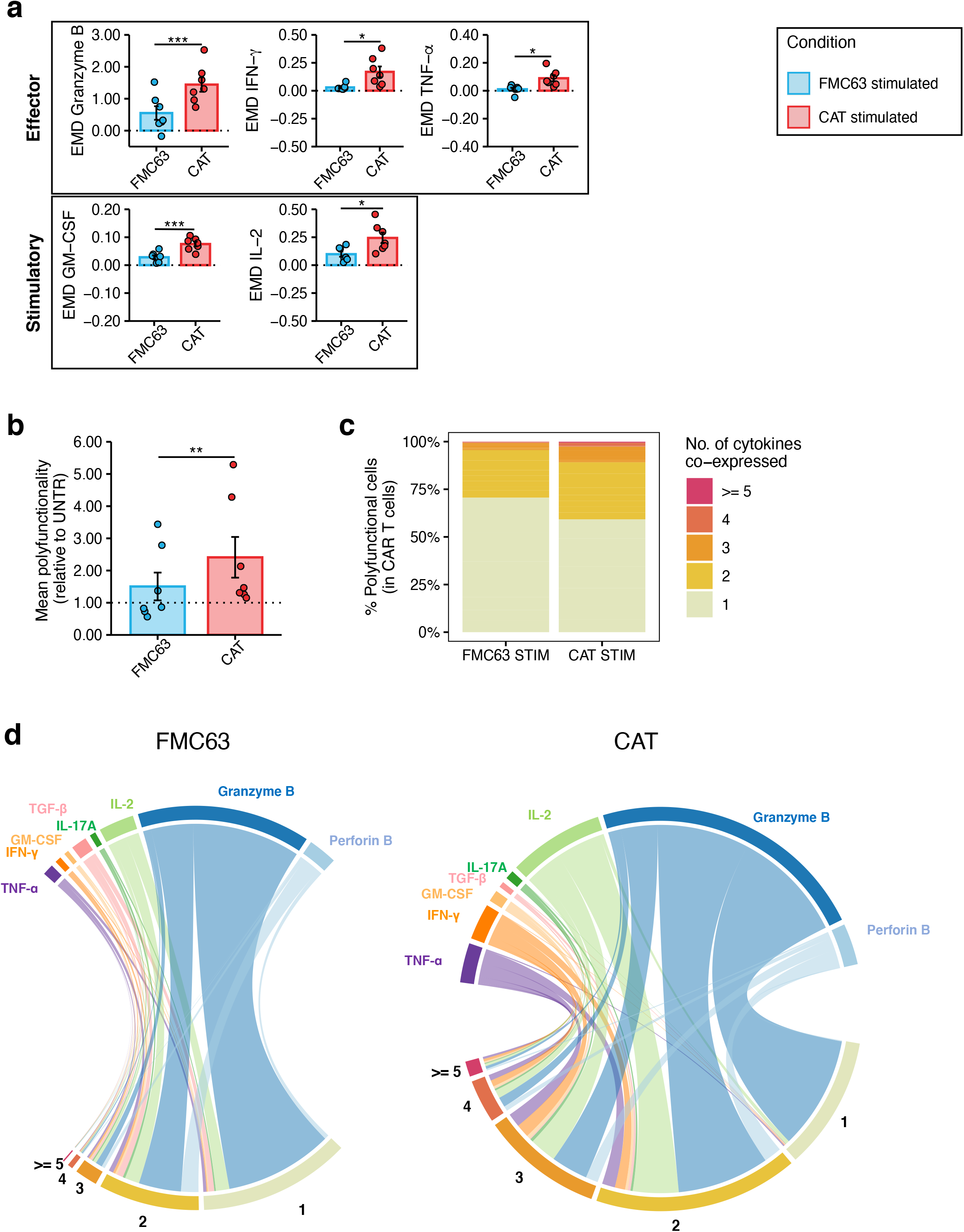
Cytokine polyfunctionality in stimulated CAR-transduced T-cells. **a** The barplots show the expression of mass cytometry EMD scores for effector (Granzyme B, IFN-γ, TNF-α) and stimulatory (GM-CSF, IL-2) cytokines in stimulated CAR T-cells. The data shown are normalized to stimulated CD3+ UNTR T-cells. The dotted horizontal line (0) represents the expression of a specific marker in stimulated CD3+ UNTR T-cells. (n = 7 HDs, n = 2 independent experiments). **b** Barplots showing the mean cytokine polyfunctionality in stimulated CAR T-cells, normalized to stimulated CD3+ UNTR T-cells. The dotted horizontal line (1) represents the mean polyfunctionality in stimulated CD3+ UNTR T-cells. (n = 7 HDs, n = 2 independent experiments). **c** The stacked barplots show the percentage of stimulated CAR T cells (CD3+mCherry+) expressing 1 to 4 or >=5 cytokines/cell as measured by mass cytometry. **d** The circos plots show all the combinations of the 8 cytokines analysed in stimulated FMC63 (left) and CAT (right) CAR T-cells by mass cytometry. The numbers indicate patterns of cytokine co-expression (from 1 to 4 or >=5 cytokines/cell). A specific colour code has been assigned to each cytokine. **a-b** Data represent mean ± se. Statistical significance was calculated by Paired t-test. *P < 0.05, ***P < 0.001. Each experimental condition is indicated by a specific colour code (FMC63 blue, CAT red).

Next, we investigated the overall pattern of cytokine co-expression in CAT and FMC63 CAR T-cell responses. The ability of a single T-cell to express simultaneously more than one cytokine (polyfunctionality) has been linked to productive immune responses^35,36^ and more recently described as a distinctive feature of CAR T-cells associated to their potency and anti-tumour efficacy ^37–39^. We measured the frequency at which the eight cytokines included in our analysis were co-expressed in single CAR T-cells, thus providing a comprehensive profile of their cytokine polyfunctionality. Upon stimulation with NALM6, 15.02% of FMC63 and 29.60% of CAT CAR T-cells were polyfunctional (expressing two or more cytokines per cell) (Supplementary Table 5n).

Furthermore, not only the frequency of polyfunctional CAR T-cells, but also the number of cytokines co-expressed was higher in CAT than in FMC63, with a stastically significant increase in their mean polyfunctionality (Fig. 4b) and a marked increment in the proportion of cells expressing combinations of three or more cytokines (2.28% in FMC63 *vs* 7.89% in CAT) (Fig. 4c and Supplementary Table 5o). The polyfunctional profiles of stimulated CAT CAR T-cell products were dominated by combinations of cytokines involving the effector molecules IFN-g, TNF-α, IL-2 and Granzyme B (Fig. 4d).

### Low-affinity CD19 CAT CAR activation priming is associated with and driven by residual CD19-expressing B-cells

To investigate whether the mechanism behind the activation priming observed in unstimulated CAT CAR T-cells was antigen-dependent or -independent, we checked if residual CD19+ B-lymphocytes were detectable in the CAR T-cell product and could serve as a potential source of antigen specific activation.

We monitored the proportion of CD19+ B-lymphocytes in culture at different timepoints. While B-cells were detectable at day 0 in all samples (5.96% of cells on average), at day 8 (5 day stimulation with CD3/CD28 beads + 3 day rest) they could only be detected in the UNTR condition (2.33% of cells on average), and had been completely depleted from both CAR constructs, as shown by FlowSOM analysis of mass cytometry data, clustering single-cells by cell types^40^, and the relative frequencies (Fig. 5a and Supplementary Fig. 7a). To test whether the activation priming observed in CAT CAR T-cells was dependent on donor B-cells, we applied the experimental setup described in Fig. 1a, only now including as additional experimental condition CAR T-cells generated from CD19-depleted PBMCs. We confirmed effective CD19 depletion by flow cytometry, with an average of residual B cells of 0.041% upon depletion as compared to 5.96% with the standard protocol (Supplementary Fig. 7b). The transduction levels obtained in the CD19-depleted CAT and FMC63 CAR T-cells were comparable based on the expression of the fluorescent reporter (mCherry) and the VCN (Supplementary Fig. 7c-d). The baseline CD4:CD8 ratios only showed a slight increase in CAT as compared to FMC63 CAR T cells (Supplementary Fig. 7e). Of note, CD19-depletion impacted on the T memory subsets composition, with CAT CAR T-cells no longer displaying any statistical difference when compared to the UNTR control, while FMC63 CAR T-cells still showing a significant increase in the proportion of T_EM_ when compared to the UNTR control (Fig.5b).

**Fig. 5:**
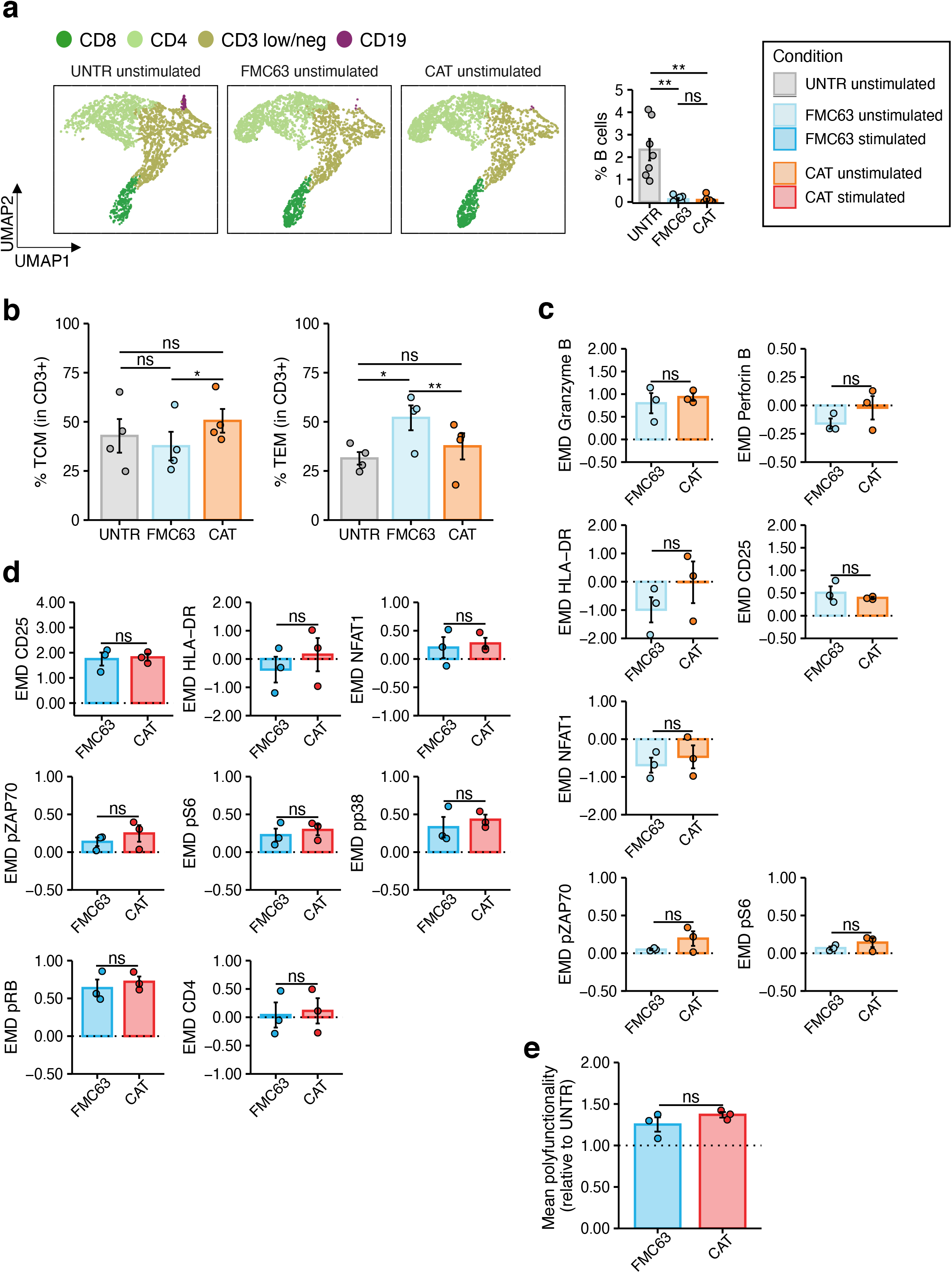
Molecular characterisation of CAR T-cells generated from CD19-depleted PBMCs. **a** (left) UMAP representation of the 4 cell populations (CD8, CD4, CD3 low/neg, CD19) identified by FlowSOM analysis in 4 representative unstimulated samples analysed by mass cytometry at 24 h post-stimulation (n = 4 HDs, n = 1 independent experiment). The cell types are indicated by different colours. (right) Percentage of residual B-cells detected by mass cytometry in unstimulated samples at 24 h post-stimulation. (n = 7 HDs, n = 2 independent experiments). **b**(left) Barplots showing the percentage of TCM (CD45RA-CD62L+) and (right) TEM (CD45RA-CD62L-) in unstimulated CD19-depleted CD3+ UNTR T-cells and FMC63 and CAT CAR T-cells measured by FACS (n = 4 HDs, n = 1 independent experiment). **c** The barplots show the expression of mass cytometry EMD scores for Granzyme B, Perforin B, HLA-DR, CD25, NFAT1, pZAP70 and pS6 in unstimulated CD19-depleted CAR T-cells at 24 h upon stimulation. The data shown are normalized to stimulated CD3+ UNTR T-cells. The dotted horizontal line (0) represents the expression of a specific marker in unstimulated CD3+ CD19-depleted UNTR T-cells. (n = 3 HDs, n = 1 independent experiment). **d** The barplots show the expression of mass cytometry EMD scores for CD25, HLA-DR, NFAT1, pZAP70, pS6, pp38, pRB and CD4 in stimulated CD19-depleted CAR T-cells at 24 h upon stimulation. The data shown are normalized to stimulated CD3+ UNTR T-cells. The dotted horizontal line (0) represents the expression of a specific marker in stimulated CD19-depleted CD3+ UNTR T-cells. (n = 3 HDs, n = 1 independent experiment). **e** Barplots showing the mean polyfunctionality in CD19-depleted CAR T-cells at 24 h upon stimulation. The data shown are normalized to stimulated CD3+ UNTR T-cells. The dotted horizontal line (1) represents the mean polyfunctionality in stimulated CD3+ CD19-depleted UNTR T-cells. (n = 3 HDs, n = 1 independent experiment). **a-e** Each experimental condition is indicated by a specific colour code (Unstimulated conditions = UNTR light grey, FMC63 light blue, CAT orange; stimulated conditions = FMC63 blue, CAT red). Barplots show mean ± se. Statistical significance was calculated by Paired t-test. *P < 0.05, **P < 0.01.

Mass cytometry analyses revealed that the residual B cells in the CAR T-cell manufacture were responsible for the activation priming observed in unstimulated CAT and FMC63 CAR T-cells (Supplementary Fig. 8, 9). While CD19 depletion led to a general decrease in the activation priming previously observed in unstimulated CAR T-cells, this reduction was more pronounced in CAT (Supplementary Fig. 8) than in FMC63 (Supplementary Fig. 9). As a result, upon B cell depletion we no longer detected statistically significant differences in the expression of T-cell activation markers (HLA-DR, CD25 and NFAT1), pro-inflammatory cytokines (Granzyme B, Perforin B) and CAR-downstream signaling molecules (pZAP70 and pS6) between CAT and FMC63 (Fig.5c). Differential protein expression between CD19-depleted CAT and FMC63 CAR T-cells was only observed for pRB and CD69 (Supplementary Fig. 7f). Both CAT and FMC63 CAR T cells exhibited increased expression of activation and cytotoxic markers with respect to the UNTR controls they are normalized to, indicating similar levels of antigen-independent activation (Fig. 5c).

When later exposed to CD19-expressing NALM6, both CD19-depleted CAT and FMC63 were able to activate, as shown by the upregulation of the expression of cytotoxic markers and cytokines compared to their unstimulated counterparts (Supplementary Fig. 10, 11). No differences in the CD4:CD8 ratios and in the expression of exhaustion markers were observed between the two stimulated CAR conditions upon CD19-depletion (Supplementary Fig. 7g, h). Most importantly, upon antigenic stimulation, CD19-depleted CAT CAR T-cells activated a molecular response with no stastically significant differences when compared to FMC63 (Fig. 5d and Supplementary Fig.7i), except for an increased expression of IL-2 (Supplementary Fig.7i). In agreement, the increased cytokine polyfunctionality observed in CAT *vs* FMC63 CAR T-cells in standard manufacture condition (Fig. 4) was no longer observed in CD19-depleted manufacture condition (Fig. 5e).

Altogether these results demonstrate that residual B-cells in the CAR T-cell manufacture can mediate an antigen dependent activation priming, which is more pronounced in low affinity CAT CAR T-cells when compared to FMC63 CAR T-cells. Such activation priming contributes to boosting CAT CAR T-cell response, as CAT CAR T-cells generated from CD19-depleted PBMCs not only do not display increased activation priming but also do not exhibit increased molecular responsed to antigenic stimulation with NALM6.

## Discussion

Modulating CAR T-cell affinity may enable us to enhance anti-tumour response and long-term tumour surveillance, while minimizing CAR T-cell related toxicity. To this aim, characterizing the molecular determinants of treatment efficacy in the low-affinity CAT CAR may inform future design and optimize their manufacture. We thus investigated the transcriptomic and proteomic phenotype of CD19 CAT CAR T-cells, compared with the widely used FMC63, to begin unravelling the molecular mechanisms behind the observed preclinical and clinical differences between these two CD19 CARs^11^.

We found that CAT CARs induce stronger activation responses than FMC63, which could be explained by the faster target off-rate of low-affinity CARs. Faster dissociation requires fewer targets to serially trigger a larger number of CARs and amplify anti-tumoural response^13^. This would align with the proposed model of temporal and spatial summation of T-cell activation, in which signals from serially triggered immunoreceptors can be accumulated and integrated overtime to reach the threshold required for T-cell activation^41^. This model may also explain why low levels of residual donor B cells in the manufacture can induce stronger activation priming in CAT CAR T cells than in FMC63. However, as CAT and FMC63 have been predicted to bind overlapping but not necessarily identical epitopes on CD19, we cannot exclude that the epitope location could also contribute, in part, to the functional differences observed^11,42^.

Our transcriptomic and protein profiling revealed that unstimulated CAT CAR T-cells are functionally closer to antigen activated CAR T-cells than FMC63 CAR T-cells, with a number of upregulated activation genes (*HLA-DBP1, HLA-DRA* etc) and proteins (HLA-DR, CD25 and NFAT1). Despite these genes/proteins being commonly associated to T-cell activation, their basal expression is also increased in memory T-cells as compared to T_NAÏVE_ cells. This is because memory T-cells are characterized by an open chromatin conformation favouring the access of transcription factors to immune response genes, thus ensuring a pool of readily available mRNAs that can be rapidly translated following stimulation^43^. This hypothesis would be in agreement with the increased proportion of T_CM_ observed in CAT CAR T cells as compared to FMC63.

By performing B cell depletion prior to CAR T-cell manufacture, we demonstrate that residual B-cells expressing CD19 in the CAR T-cell product are responsible for the activation priming observed in CAR T-cells, which was more pronounced in CAT when compared to FMC63. This is consistent with the hypothesis that CAT serial triggering may amplify T-cell activation from lower antigen levels^13^ and points to a boosting role of low dose CD19-priming during CAR T-cell manufacture. Recent results have shown that antigen-independent induction of CAR T-cell priming, by either 4-1BB-based tonic signalling^22^ or by low-dose of hypomethylating agents^23^ can lead to enhanced CAR T anti-tumour functions *in vivo*. Our results indicate that low dose antigen-specific priming can also promote CAR T-cell functionality in a CAR construct specific manner with an enhanced effect in low-affinity CAR T-cells. This results may appear controversial, relatively to the increasing evidence that CD4/CD8 T-cell positive selection prior to manufacture leads to CAR T-cell products outperforming those derived from unfractionated leukaphereses. However, the improved performance of CD4/CD8 CAR T cell manufactures has been mechanistically linked to the depletion of myeloid and NK cells negatively impacting CAR T cell transduction, activation and expansion, while the role of B cells has not been extensively investigated in this setting^44–46^.

When stimulated with CD19-expressing NALM6 cell line, CAT CAR T-cells have a distinct transcriptomic and protein response to CD19 antigenic stimulation from FMC63 CAR T-cells with increased expression of proliferation, activation and cytotoxic markers at both RNA and protein levels. *In vivo* clonal kinetics analyses have shown that single CD19 CAR T-cells with higher expression of cytotoxic related genes in the manufacture product, many of which in common with ours (including *IFNG, HLA-DRA, CCL4*), gave rise to superior *in vivo* expansion and survival, significantly contributing to later timepoints after adoptive transfer in patients^19,31^. Analysis of gene expression also revealed the upregulation of chemoattractive cytokines CCL4 and CCL3L1 in CAT, which induce T-cell homotypic interactions and promote the reciprocal exchange of self-reinforcing signals such as OX40L,^26,29^ which are upregulated in CAT as well. Single-cell transcriptomic studies have recently revealed that subsets of CAR T with elevated expression of CCL3 and CCL4 are associated to longer persistence in vivo^31^ and achievement of complete remission^32^. Mass Cytometry analyses revealed a CAT CAR T enhanced activation protein profile, as measured by the increased expression of T-cell activation markers (CD25, HLA-DR) and CAR downstream signaling effectors (pZAP70, pp38, NFAT1, pCREB, FOXP3). Furthermore, CAT CAR T-cells showed increased mTORC signaling (pS6 and pRB), which is commonly associated to cell proliferation and protein translation. Altogether, CAT CAR T-cells distinct gene expression and protein profiles are very much in line and likely responsible for the enhanced proliferative responses that CAT CAR T-cells exhibited *in vitro*, in *in vivo* murine models and in patients, as previously reported by Ghorashian et al.^11^.

Upon stimulation, CAT CAR T-cells were also characterized by a unique polyfunctional pattern of cytokine expression, with a marked increase in the frequency of single CAR T-cells expressing >=3 cytokines when compared to FMC63. CAT polyfunctional profile was dominated by combinations of effector cytokines, consistent with their potent anti-tumour activity. It has been suggested that the ability of CAR T-cells to produce multiple cytokines (polyfunctionality) in response to antigen exposure is associated with improved anti-tumour responses *in vivo*^37^ and cytokine polyfunctionality has been recently proposed as a criteria to predict CAR T-cell potency ^39^. The increased cytokine polyfunctionality observed in CAT CAR T-cells contributes to explain their increased cytotoxic potential previously observed *in vitro settings* and in *in vivo* murine models^11^.

In conclusion, we describe the molecular mechanisms underlying the low-affinity CAT CAR T-cell functional phenotype. We provide evidence that the potent and long-term anti-tumour responses observed with low-affinity CAT CAR T-cells^11^ reflect a distinct pattern of both activation priming and cytokine polyfunctionality. We show that low affinity CAT CAR T-cells are preferentially primed by low concentration of CD19-expressing B-cells present in the manufacture and such priming is instrumental to their higher cytotoxic response upon stimulation. Although our observations are limited to one low-affinity CAR, future work extending this characterization to a panel of low-affinity CARs, may reveal at which extent these findings are generalisable. Overall, our work has important implications for the future design of versatile CAR T-cells manufacture protocols, capable of boosting efficacy and long-term persistence.

## Methods

### Human peripheral blood samples and cell lines

Peripheral blood was collected from 20 HDs after having obtained informed consent. Fresh mononuclear cells were isolated by Ficoll density gradient centrifugation (day 0) and used for experiments (standard condition). CD19+ B cells were magnetically removed from freshly isolated PBMCs using CD19 Microbeads (Miltenyi Biotec, Bergisch Gladbach, Germany) according to manufacturer’s recommendations (depleted condition). All analyses were performed on fresh T- or CAR T-cells except for one 96 h memory subset FACS analysis. HD details and relative experimental procedure are reported in Supplementary Table 1. The human ALL cell line NALM6 was purchased from DSMZ. Cells were cultured according to manufacturer’s instructions in complete RPMI 1640 medium, GlutaMAX™ Supplement (Gibco™, Thermo Fisher Scientific, Waltham, MA, USA) supplemented with 10% of heat-inactivated Foetal Bovine Serum (FBS) (Gibco™, Thermo Fisher Scientific), 100 IU penicillin and 100 μg/ml streptomycin (Sigma-Aldrich, St. Louis, MO, USA).

### *Ex vivo* T-cell expansion

Freshly isolated PBMCs and CD19-depleted PBMCs were cultured in TexMACS™ medium (Miltenyi Biotec, Bergisch Gladbach, Germany), an optimised T-cell medium. To induce T-cell expansion, CD3/CD28 beads (CTS™ Dynabeads™ CD3/CD28, Thermo Fisher Scientific) were added to cells in MACS GMP Cell Differentiation Bags (Miltenyi Biotec) at a 1:3 lymphocyte:bead ratio. CD3/CD28 beads were magnetically removed from the culture on day 5 and CAR T-cells rested for 48 h before proceeding to antigen stimulation.

### CAR construct generation

CD19 FMC63 and CAT CAR transfer vectors were generated as previously described^11^ and were provided by P.J.A.. The vector structure is schematised in Supplementary Fig. 1a.

### Lentiviral production by transient transfection

Lentiviral production was performed by transient transfection according to the following procedure. The day before transfection, 5.5 x 10^6^ 293T cells were seeded per 175 cm^2^ flasks in complete DMEM, high glucose, GlutaMAX™ Supplement (Gibco™, Thermo Fisher Scientific) supplemented with 10% FBS.

To generate viral particles, 293T cells were co-transfected with second generation LV packaging plasmids pMD2.G and pCMV-dR8.74 (plasmidFactory, Bielefeld, Germany) and CAR transfer vectors using GeneJuice^®^ Transfection Reagent (Merck-KGaA, Darmstadt, Germany) according to manufacturer’s recommendations. LV supernatants were collected at 48 and 72 h post-transfection, pooled, filtered and stored at −80°C before use.

### Lentiviral transduction of T-cells

Following overnight activation with CD3/CD28 beads, 0.5 x 10^6^ beads-activated T-cells were suspended in 0.5 ml of TexMACS, transduced to express CD19 CAR construct (FMC63 or CAT) with 1 ml of LV supernatant in RetroNectin®(Takara Bio, Kusatsu, Shiga, Japan)-coated 24-well plates and spinoculated at 1,000 g for 40 min at room temperature. Generally, 2-10 x 10^6^ beads-activated T-cells per donor per construct were seeded for transduction. Transduction efficiency was assessed by FACS evaluating mCherry (LV fluorescent marker) expression, at day 8 (7 days after transduction). The copies of lentiviral vectors integrated in the human genome (vector copy number, VCN) were quantified at day 11 (10 days after transduction) using a TaqMan real-time PCR assay targeting the woodchuck hepatitis virus post-transcriptional regulatory element (WPRE) in the LV construct and the human beta-actin gene (Integrated DNA Technologies, Coralville, Iova, USA).

UNTR beads-activated T-cells per each donor were used as a control throughout the experiment and were subjected to the same experimental conditions as LV-transduced ones.

### NALM6 and CAR T-cells co-cultures

To assess the anti-tumour activity of CD19 CAR T-cells, we set-up co-cultures with CD19-expressing NALM6 for 24 or 96 h (stimulated conditions) on day 7 (6 days after transduction and 2 days post-CD3/CD28 beads removal). Briefly, beads-activated T-cells were incubated for 48 h in optimised T-cell medium before stimulation. 0.1 x 10^6^ UNTR or FMC63 or CAT beads-activated T-cells were seeded in 96-well plates in complete TexMACS medium in a 1:1 ratio with irradiated (40 Gy) NALM6. Generally, 0.1-6 x 10^6^ beads-activated T-cells per donor per condition (UNTR, FMC63 and CAT) were stimulated. Unstimulated, beads-activated T-cells per each donor, condition and time point were kept in culture under the same experimental conditions as antigen-stimulated T-cells.

### Flow cytometry activated cell sorting and antibody staining

All FACS experiments included Fluorescence minus one (FMO) and single-antibody stained BD™CompBeads (BD Biosciences) controls to set expression threshold and to calculate compensation, respectively. Live cells were selected based on their non-permeability and lack of fluorescence associated with DAPI Solution (BD Biosciences, Franklin Lakes, NJ, USA) or 7-AAD Viability Staining Solution (BioLegend, San Diego, CA, USA). The combination of monoclonal antibodies used to identify T-cell subsets is described in the main text. Following 24 or 96 h of stimulation, the phenotype of live T-/CAR T-lymphocytes in all the experimental conditions was assessed using the following antibodies: BV510-anti CD3 (clone OKT3, BioLegend), APC-CY7-anti CD3 (clone UCHT1, BioLegend), APC-CY7-anti CD4, FITC-anti CD4 (clone SK3, BioLegend), PerCP/CY5.5-anti CD8, BV510-anti CD8 (clone RPA-T8, BioLegend), AF647-anti CD62L, APC-anti CD62L (clone DREG-56, BioLegend), BV785-anti CD45RA (clone HI100, BioLegend), V450-anti CD45RA (clone HI100, BD Biosciences), AF647-anti TIM3 (clone 7D3, BD Biosciences), BB515-anti PD1 (clone EH12.1, BD Biosciences), BV605-anti LAG3 (clone 11C3C65, BioLegend).

PBMCs were labelled with PE-anti CD19 (clone HIB19; BioLegend) to detect B-cells in the samples soon after isolation and to check purity after CD19-depletion.

For CAR staining, CAR T-cells were incubated with CD19 CAR Detection Reagent human, Biotin (Miltenyi Biotec) followed by incubation with APC-anti biotin Antibody, REAfinity™ (Miltenyi Biotec) and BV510-anti CD3 (clone OKT3; BioLegend) according to manufacturer’s recommendations.

Experiments were performed on a cell sorter FACSAria™ III (BD Biosciences) and on a CytoFLEX analyzer (Beckman Coulter Inc., Brea, CA, USA) and analysed with FlowJo™ software v10.6.1 (BD Biosciences) and Cytobank platform (www.cytobank.org).

### RNA sequencing libraries preparation and sequencing

RNA sequencing libraries were generated from 8 donors (HD1-HD8), across six experimental conditions, UNTR T-cells or FMC63 or CAT CAR T-cells in either absence or presence of NALM6 (unstimulated or stimulated). 1,000 T-cells/CAR T-cells were FACS-sorted in single PCR tubes containing 4 ul of QIAGEN lysis buffer from QIAseq FX Single Cell RNA Library Kit (QIAGEN, Hilden, Germany) and stored at −80°C. The kit steps include reverse transcription, cDNA amplification and PCR-free NGS library preparation. Libraries were purified using AMPure XP magnetic beads (Beckman Coulter Inc.,) according to the manufacturer instructions. After beads purification the libraries were resuspended in 50 μL of buffer EB (Qiagen) and stored at −20 °C. Size and concentration of the libraries generated was assessed using High Sensitivity Bioanalyzer (Agilent, Santa Clara, CA, USA) and Qubit High-Sensitivity DNA kit (Invitrogen™, Thermo Fisher Scientific), respectively. Libraries were equimolarly pooled to a final concentration of 10 nM and were sequenced with Illumina HiSeq 3000 (150 bp pair-end reads) at UCL Genomics.

### RNA sequencing analysis

Sequencing quality control and mapping was performed on Fedora 32. DGE analysis and downstream analyses were performed in *R 4.0.2* on Windows 10 (build 18362). Additional details can be found in the transcriptomic Supplementary Bioinformatics Material (SBM). An FDR smaller than 0.1 (when multiple tests were performed) or a *P* value smaller than 0.05 (when only one test was performed) was considered statistically significant. Quality control and adapters trimming was performed with *FastQC 0.11.9^47^* and *cutadapt 2.10^48^*. Sequences were mapped with *STAR 2.7.3a^49^* to the release 31 of the Human Genome (GRCh38.p12)^50^ and summarised with *featureCounts* from the *subread 2.0.0* software package^51^. Only genes annotated as protein-coding genes were included into the DE and downstream analyses. These were further filtered by the *filterByExpr* function from the *edgeR* package ^52^ in order to remove lowly expressed genes. DE was assessed with *DESeq2^53^*. FDR were calculated using Benjamini-Hochberg correction^54^.

Only the genes which passed the *filterByExpr* filtering were used for GSEA pathway analysis. GSEA was performed with *fgsea* package^55^ on a subset of pathways from the MSigDB’s Hallmarks pathways^56^ (see SBM Table III). The log2Fold Change of the DE conditions compared with their matched UNTR condition was used as an input of the analysis. Relative cell type enrichment was performed with *xCell^57^* on a subset of cell types (SBM Table IV).

### Mass cytometry analysis

Following 24 h of stimulation, samples in all experimental conditions were treated with Brefeldin A (BioLegend) (1:1,000) at 37°C for 4 h to favour intracellular cytokine accumulation and were fixed with 1.6% of formaldehyde for 10 min at room temperature. Fixed samples were stained with 0.05 μM Cell-ID™ Cisplatin-194-Pt (Fluidigm, South San Francisco, CA, USA), a live-dead marker, for 1 min on a rocker. Samples for all experimental conditions from a given donor were then barcoded using the Cell-ID™ 20-Plex Pd Barcoding Kit (Fluidigm) according to manufacturer’s instructions. 12-16 samples were labelled concurrently in the same tube following a three-step staining protocol. Briefly, 2.6-5 x 10^6^ cells were firstly stained with antibodies against surface markers (CD3, CD4, CD8a, CD19, CD22, CD25, CD44, CD45RA, CD62L, CD69, CD127, HLA-DR and TIM-3). Following permeabilization with eBioscience™ Foxp3/Transcription Factor Staining Buffer Set (Thermo Fisher Scientific) for 30 min at 4°C, cells were subsequently stained for the detection of intracellular cytokines (IL-4, IL-5, IL-6, IL-23, IL-17A, IL-2, TNF-α, IFN-γ, GM-CSF, TGF-β, Granzyme B and Perforin B) and cytoplasmic protein (RFP and mCherry, used to detect mCherry expression). Cells were then fixed in ice-cold 50% Methanol for 10 min at 4°C and underwent antibody staining for nuclear targets (cleaved Caspase 3, Cyclin B1, FOXP3 and NFAT1) and phosphoproteins (p4E-BP1, pAMPKa, pBAD, pBTK, pCREB, pERK1/2, pHistone H2A.X, pHistone H3, pMKK3/MKK6, pp38, pPDPK1, pRB, pS6, pSRC, pSTAT4, pSTAT5 and pZAP70). Two independent mass cytometry experiments were performed and the antibodies used are listed in Supplementary Table 3. Stained cells were fixed with 1.6% of formaldehyde for 10 min at room temperature on a rocker and incubated with Cell-ID™ Intercalator-Ir (Fluidigm) overnight at 4°C. The following day, experiments were performed on a Helios mass cytometer (Fluidigm). EQ™ Four Element Calibration Beads (Fluidigm) were added to cell suspensions immediately before acquisition to guarantee inter-sample comparability.

### Mass cytometry data pre-processing and analysis

The resulting FCS files were normalised against EQ™ Four Element Calibration Beads, debarcoded and uploaded on Cytobank platform. Viable single-cells were cleaned based on Gaussian parameters (Event length, Centre, Offset and Residual), DNA content (193-Ir DNA) and Cisplatin (194-Pt) and gated on CD19, CD3 and mCherry markers to detect B-cells and T-/CAR T-lymphocytes. Cell populations were exported from Cytobank and computationally analysed in Python (Python version 3.7.6) or in R (R version 3.6.1) as described below.

Specifically, EMD^17^ scores used for PCA were computed separately for each stimulation condition between CAR T-lymphocytes and UNTR T-lymphocytes and the entire population as previously described^58^ using CyGNAL repository (<u>10.5281/zenodo.4587193</u>).

The PCA performed on EMD scores of all 32 markers and the event counts in CD3+ T-cells from HD4-HD8 across all the experimental conditions highlighted the lack of activation of HD6 CAR T-cells upon antigen stimulation (data not shown). For this reason, HD6 was excluded from the analyses. PCA in Fig. 2b was generated on EMD scores of all 32 markers calculated on CD3+ T-cells across all experimental conditions at 24 h post-stimulation.

EMD^17^ scores in Fig. 2e, 3e, 4a, 5c-d, Supplementary Fig. 3c, 5-10 were computed separately for each given donor, stimulation condition and manufacture condition between CAR T-lymphocytes and the corresponding UNTR T-cell population.

For the comparison between CAR T-cell constructs (Fig. 2e, 3e, 4 and Supplementary Fig. 3c) and between stimulation conditions (Supplementary Fig. 5 and 6) in the standard manufacture condition, only the markers targeted with the same mass cytometry antibody between the two experiments were investigated.

Additionally, pre-gated FCS files were imported in R, arcsinh transformed (cofactor = 5) and cell clustered using the R packages *FlowSOM^40^* and *ConsensusClusterPlus^59^* as computed in^60^. The code for the dimensional reduction analysis provided^60^ was modified using R package *umap* for UMAP analysis. UMAP analyses were performed downsampling at 1,000 cells and using default parameters.

Cytokine co-expression analysis was computed in R on CAR T (CD3+mCherry+) and UNTR (CD3+) T pre-gated FCS files concatenated per donor for each given experimental condition. An expression threshold of 1 was set for all the arcsinh transformed cytokine markers except for TGF-ß, for which a threshold of 4 was selected. Chord diagrams of cytokine co-expression were created with the R package *circlize^61^*.

### Statistical analyses

Data are shown as mean ±standard error (se) and statistical analyses were performed in R calculating Paired samples t-tests across experimental conditions unless otherwise stated.

EMD-related statistical analyses between selected experimental conditions and the results of polyfunctionality analysis are reported in Supplementary Table 5.

## Data availability

The RNA sequencing data and analyses are available at NCBI’s Gene Expression Omnibus (GEO) data repository with the accession code GSE157584 and in GitHub (https://github.com/EduardoGCCM/CATvsFMC63_Michelozzi).

Mass cytometry raw and processed data will be made publicly available at https://community.cytobank.org/.

---

**Supplementary Fig. 1:**
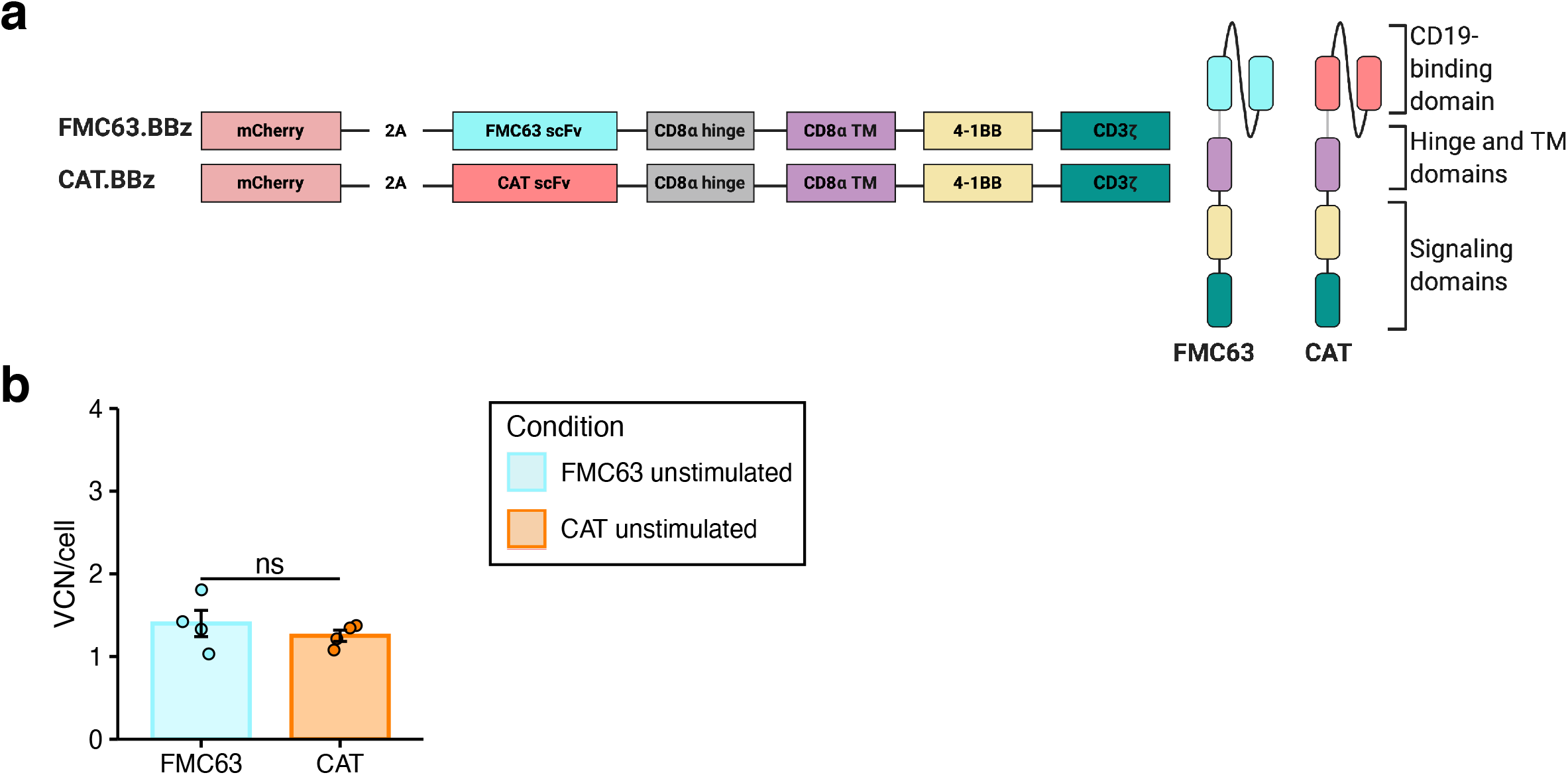
Generation and phenotypic characterisation of CAR T-cells from HD-PBMCs. **a** (left) Schematic representation of the FMC63 and CAT bicistronic lentiviral vector constructs and (right) of FMC63 and CAT CAR structure. Both constructs are second generation CARs composed by an anti-CD19 scFv, derived from FMC63 or CAT13.1E10 hybridoma, a CD8a hinge, a CD8a transmembrane domain (TM), a 4-1BB costimulatory domain and a CD3z signalling domain linked to a reporter gene (mCherry), marker of transduction. **b** The barplots show the vector copy number (VCN) quantification in unstimulated FMC63 and CAT CAR T-cells. Each dot represents an individual donor (mean of technical triplicates). Each experimental condition is indicated by a specific colour code (FMC63 light blue, CAT orange). Data represent mean ± se (n = 4 HDs, n = 1 independent experiment). Statistical significance was calculated by Paired t-test.

**Supplementary Fig. 2:**
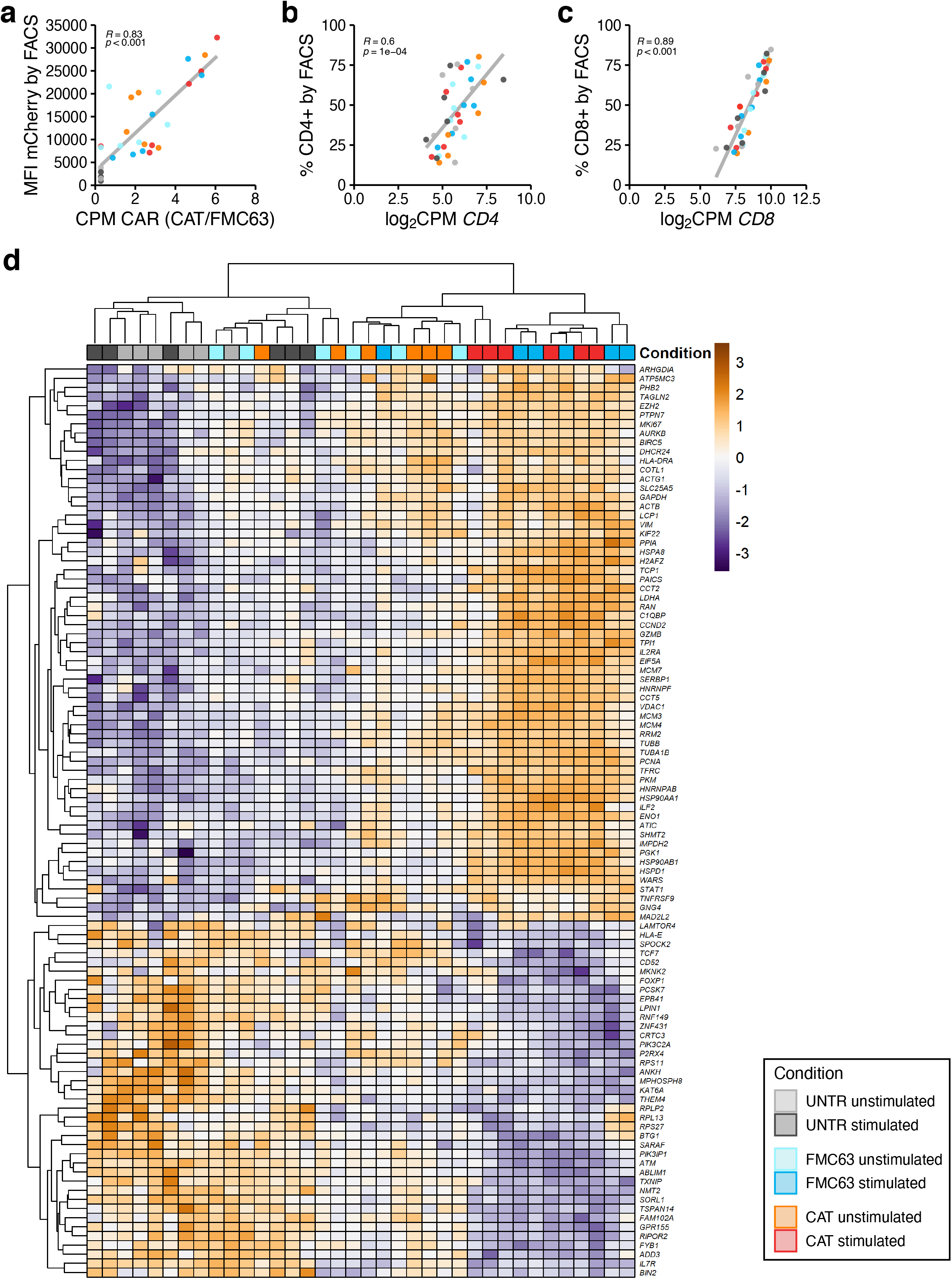
RNA-seq analyses of unstimulated and stimulated untransduced and CAR transduced T-cells. **a** Correlation of mCherry MFI measured by FACS with normalised RNA-seq counts (CPM) aligning to the scFv region of each of the two CARs (n = 6 HDs, n = 2 independent experiments). Pearson correlation and P values are displayed. **b** Correlation of the percentage of CD4+ T-cells detected by FACS with CD4 mRNA levels (n = 6 HDs, n= 2 independent experiments). Pearson correlation and P values are displayed. **c** Correlation of the percentage of CD8+ T-cells detected by FACS with CD8 mRNA levels (n=6 HDs, n = 1 independent experiment). Pearson correlation and P values are displayed. **d** Heatmap showing the hierarchical clustering of all the experimental conditions analysed displaying the top 100 variable genes from RNA-seq PC1 (n = 6 HDs, n = 2 independent experiments). **a-d** Each experimental condition is indicated by a specific colour code (Unstimulated conditions = UNTR light grey, FMC63 light blue, CAT orange; stimulated conditions = UNTR grey, FMC63 blue, CAT red).

**Supplementary Fig. 3:**
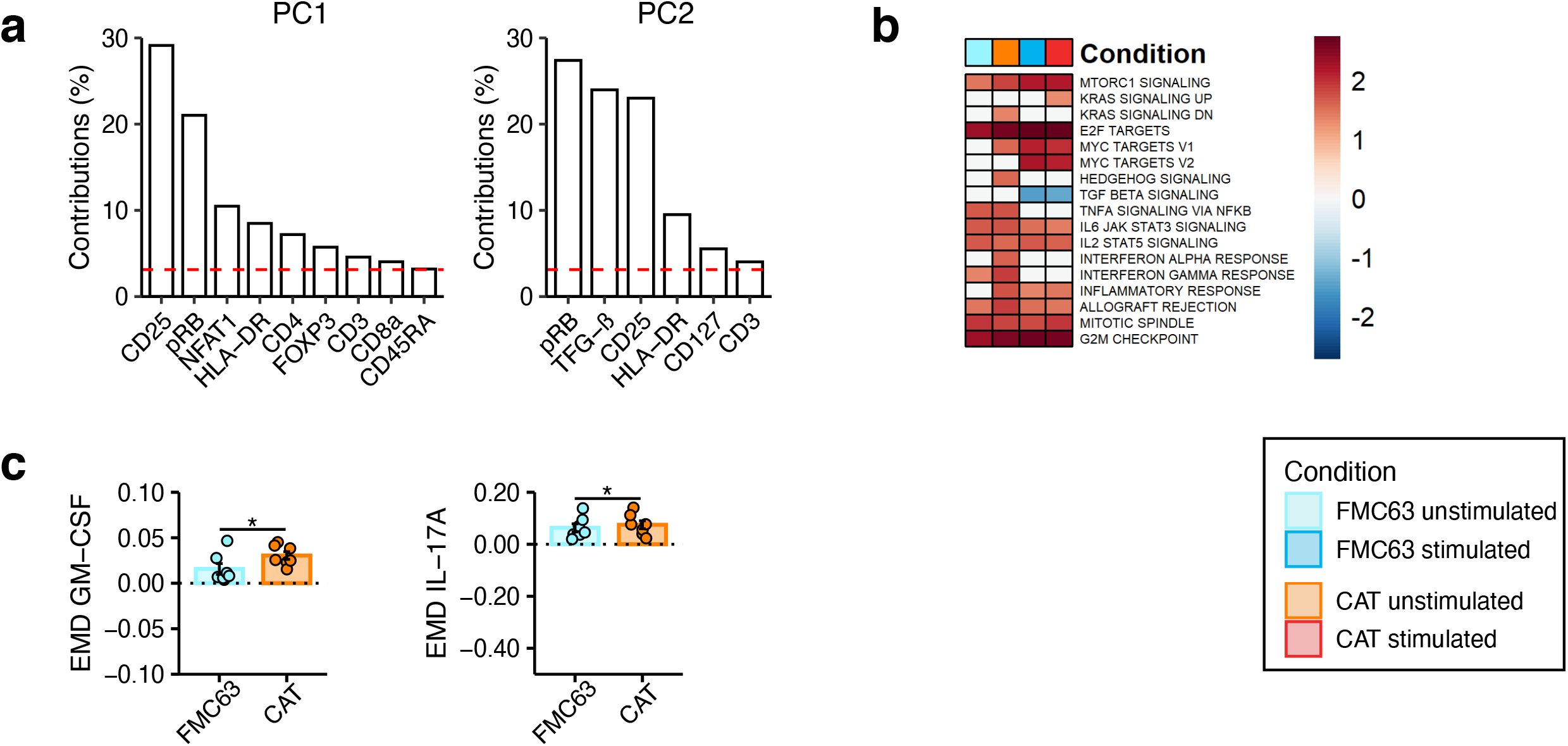
RNA-seq and mass cytometry analyses of unstimulated and stimulated untransduced and CAR-transduced T-cells. **a** Bar plots of the top contributing variables to PC1 and PC2 in Fig. 2b. The dashed horizontal red line represents the expected average contribution of the variables. **b** GSEA of HALLMARK gene sets for (1) FMC63 or (2) CAT CAR T unstimulated condition vs UNTR unstimulated cells, (3) FMC63 or (4) CAT CAR T stimulated condition vs UNTR stimulated cells (n = 6 HDs, n = 2 independent experiments). Non-significant enrichment scores were set to zero. Gene sets with no significant enrichment scores are not shown. **c** The barplots show the expression of mass cytometry EMD scores for GM-CSF and IL-17A in unstimulated CAR T-cells at 24 h upon stimulation. The dotted horizontal line (0) represents the expression of a specific marker in unstimulated CD3+ UNTR T-cells. Data represent mean ± se (n = 7 HDs, n = 2 independent experiments). Statistical significance was calculated by Paired t-test. *P < 0.05. **b-c** Each experimental condition is indicated by a specific colour code (Unstimulated conditions = FMC63 light blue, CAT orange; stimulated conditions = FMC63 blue, CAT red).

**Supplementary Fig. 4:**
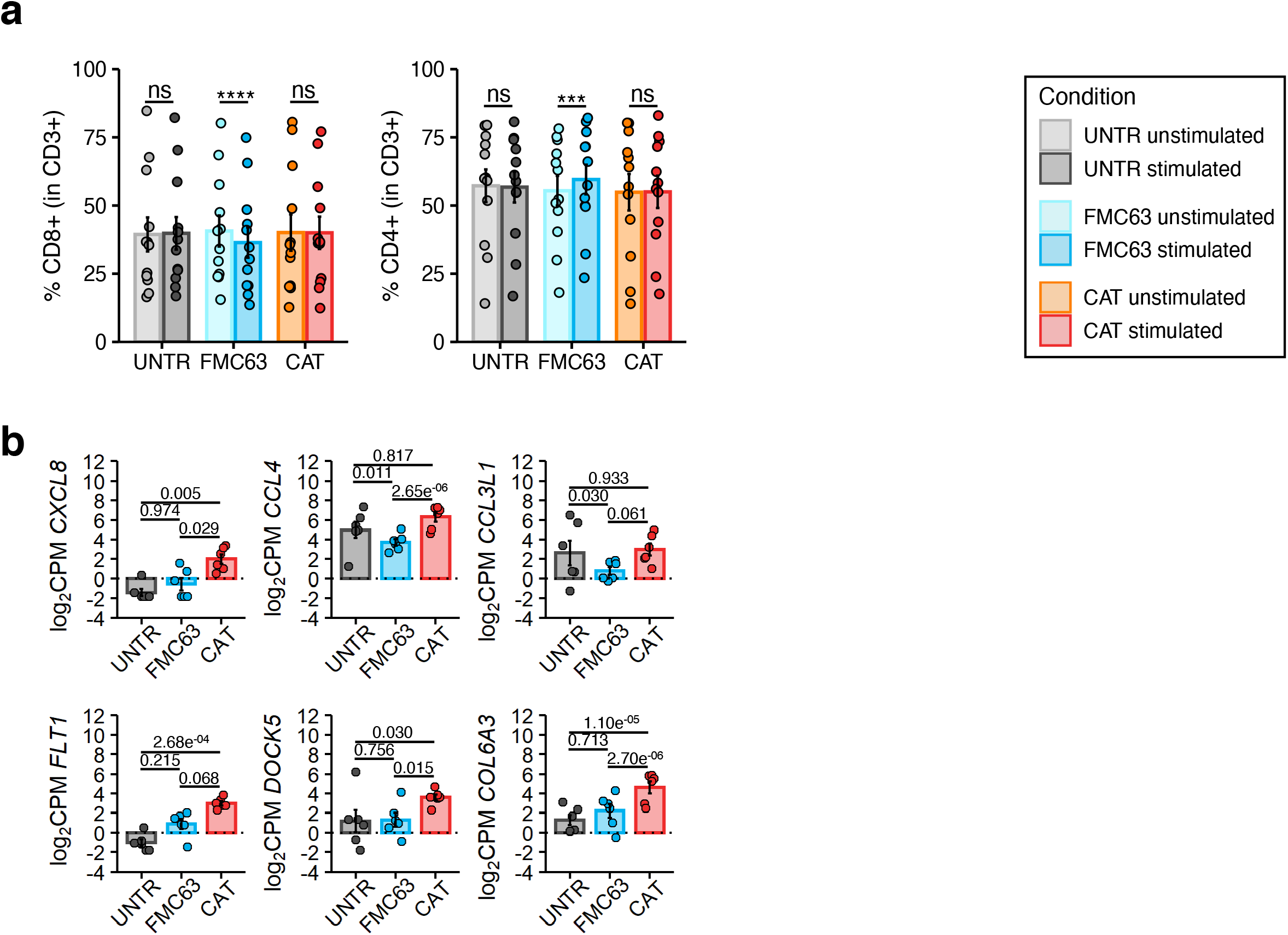
Phenotypic and molecular characterisation of stimulated CAR T-cells. **a** (left) Percentage of CD8+ cells in unstimulated and stimulated UNTR T-cells and FMC63 and CAT CAR T-cells. (right) Percentage of CD4+ cells in unstimulated and stimulated UNTR T-cells and FMC63 and CAT CAR T-cells (n = 12 HDs, n = 3 independent experiments). Statistical significance was calculated by Paired t-test. ***P < 0.001, ****P < 0.0001. **b** The barplots show the expression of selected DE genes (FDR < 0.1) in stimulated untransduced and transduced T-cells (n = 6 HDs, n = 2 independent experiments). **a-b** Barplots show mean ± se. Each experimental condition is indicated by a specific colour code (Unstimulated conditions = UNTR light grey, FMC63 light blue, CAT orange; stimulated conditions = UNTR grey, FMC63 blue, CAT red).

**Supplementary Fig. 5:**
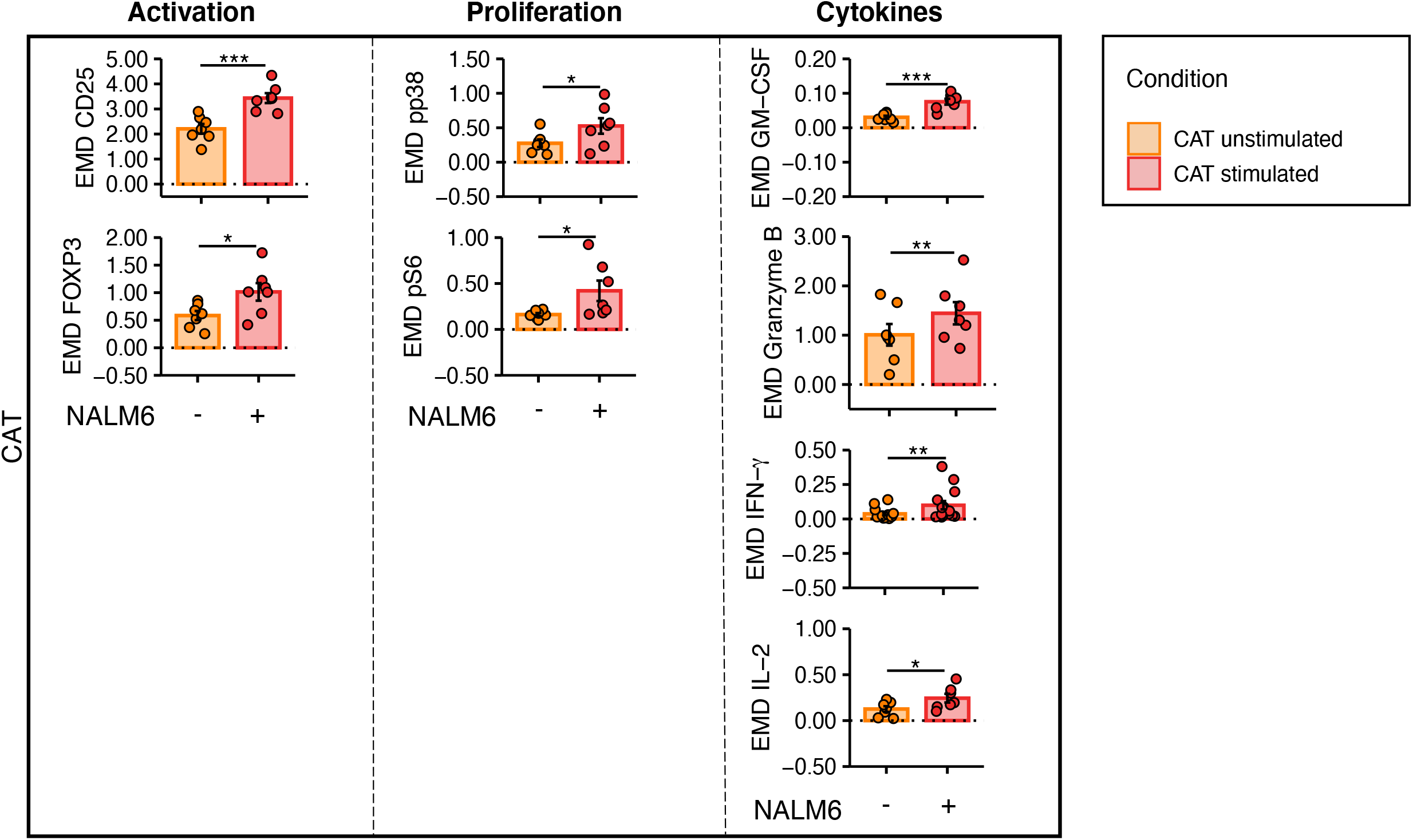
Mass cytometry EMD scores in unstimulated and stimulated CAT CAR transduced T-cells. The barplots show mass cytometry EMD scores for activation/proliferation markers and cytokines differentially expressed between unstimulated and stimulated CAT CAR T-cells. The dotted horizontal line (0) represents the expression of a specific marker in the UNTR condition, used as a reference. (n = 7 HDs, n = 2 independent experiments). Data represent mean ± se. Statistical significance was calculated by Paired t-test. *P < 0.05, **P < 0.01, ***P < 0.001. Each experimental condition is indicated by a specific colour code (Unstimulated condition orange; stimulated condition red).

**Supplementary Fig. 6:**
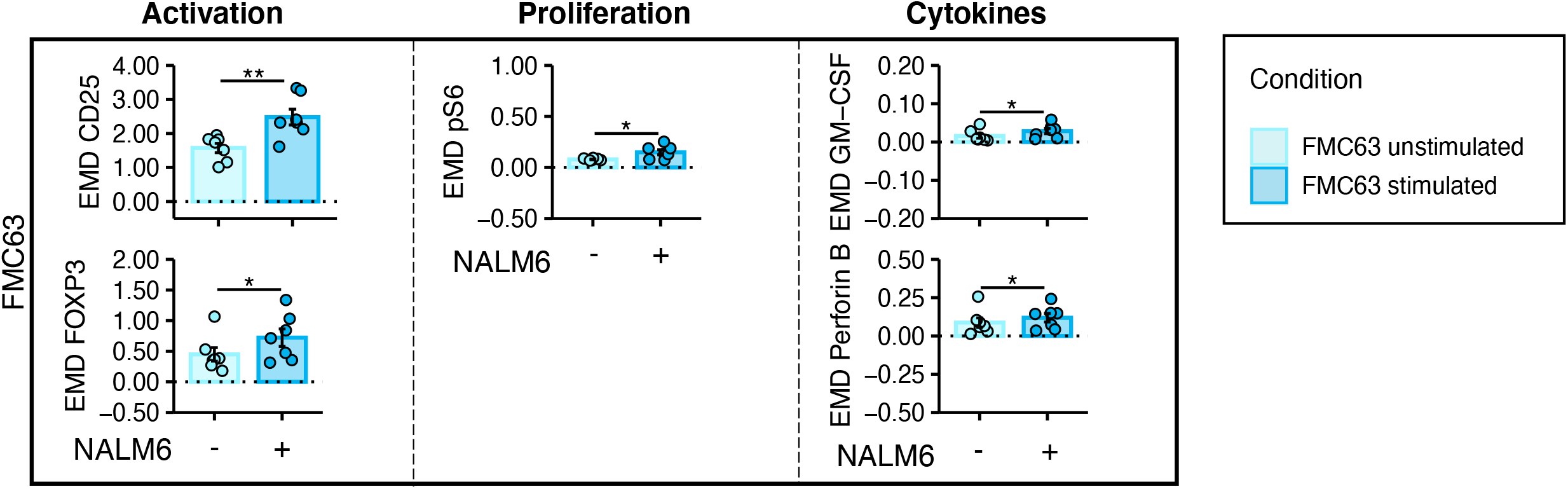
Mass cytometry EMD scores in unstimulated and stimulated FMC63 CAR transduced T-cells. The barplots show mass cytometry EMD scores for activation and proliferation markers differentially expressed between unstimulated and stimulated FMC63 CAR T-cells. The dotted horizontal line (0) represents the expression of a specific marker in the UNTR condition, used as a reference. (n = 7 HDs, n = 2 independent experiments). Data represent mean ± se. Statistical significance was calculated by Paired t-test. *P < 0.05, **P < 0.01. Each experimental condition is indicated by a specific colour code (Unstimulated condition light blue; stimulated condition blue).

**Supplementary Fig. 7:**
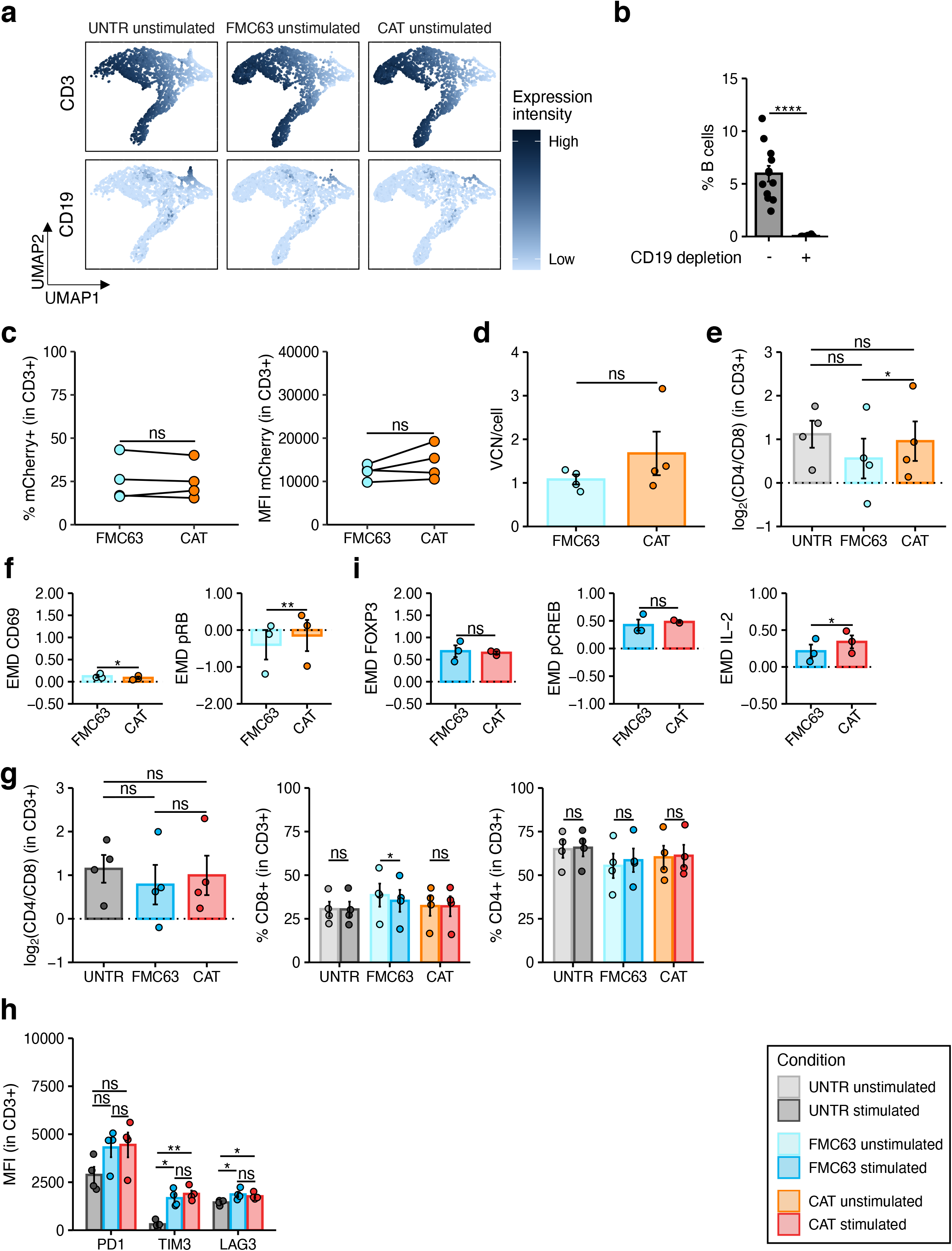
Phenotypic and molecular characterisation of untransduced and transduced T-cells derived from CD19-depleted PBMCs. **a** UMAP showing the expression of selected markers (CD3, CD19) detected by mass cytometry at 24 h post-stimulation in unstimulated CD3+ cells across all the experimental conditions (n=4 HDs, n=1 independent experiment). The colour scale reflects the protein expression levels. **b** The barplots show the percentage of B-cells detected at FACS at day 0 in freshly isolated PBMCs or in CD19-depleted PBMCs (n = 12 HDs, n = 3 independent experiments). **c** (left) Spaghetti plots showing transduction levels of CAR T-cells as percentage of mCherry+ (in CD3+) and (right) as MFI of mCherry in unstimulated transduced CD19-depleted T-cells measured by FACS at 7 days post-transduction. Lines connect results from individual donors (n= 4 HDs, n = 1 independent experiment). **d** The barplots show the integrated vector copy number (VCN) in unstimulated CD19-depleted FMC63 and CAT CAR T-cells. Each dot represents an individual donor (mean of technical duplicates) (n= 4 HDs, n = 1 independent experiment). **e** Variation (log2 fold change) of CD4 and CD8 proportion in unstimulated CD19-depleted UNTR T-cells and FMC63 and CAT CAR T-cells measured by FACS. The dotted horizontal line (0) represents the conditions in which CD4=CD8. (n=4 HDs, n=1 independent experiment). **f** The barplots show the expression of mass cytometry EMD scores for CD69 and pRB in unstimulated CD19-depleted CAR T-cells at 24 h upon stimulation. The dotted horizontal line (0) represents the expression of a specific marker in unstimulated CD3+ CD19-depleted UNTR T-cells. (n = 3 HDs, n = 1 independent experiment). **g** (left) Variation (log2 fold change) of CD4 and CD8 proportion in stimulated CD19-depleted UNTR T-cells and FMC63 and CAT CAR T-cells measured by FACS. The dotted horizontal line (0) represents the conditions in which CD4=CD8. (middle) Percentage of CD8+ cells in unstimulated and stimulated CD19-depleted UNTR T-cells and FMC63 and CAT CAR T-cells. (right) Percentage of CD4+ cells in unstimulated and stimulated CD19-depleted UNTR T-cells and FMC63 and CAT CAR T-cells (n = 4 HDs, n = 1 independent experiment). **h** Barplots showing the expression of T-cell exhaustion markers (PD1, TIM3 and LAG3) as MFI in stimulated CD19-depleted CD3+ UNTR T-cells and FMC63 and CAT CAR T-cells measured by FACS (n = 4 HDs, n = 1 independent experiment). **i** The barplots show the expression of mass cytometry EMD scores for FOXP3, pCREB and IL-2 in stimulated CD19-depleted CAR T-cells at 24 h upon stimulation. The dotted horizontal line (0) represents the expression of a specific marker in stimulated CD3+ CD19-depleted UNTR T-cells. (n = 3 HDs, n = 1 independent experiment). **b-i** Each experimental condition is indicated by a specific colour code (Unstimulated conditions = UNTR light grey, FMC63 light blue, CAT orange; stimulated conditions = UNTR grey, FMC63 blue, CAT red). Data represent mean ± se. Statistical significance was calculated by Paired t-test. *P < 0.05, **P < 0.01, ****P < 0.0001.

**Supplementary Fig. 8:**
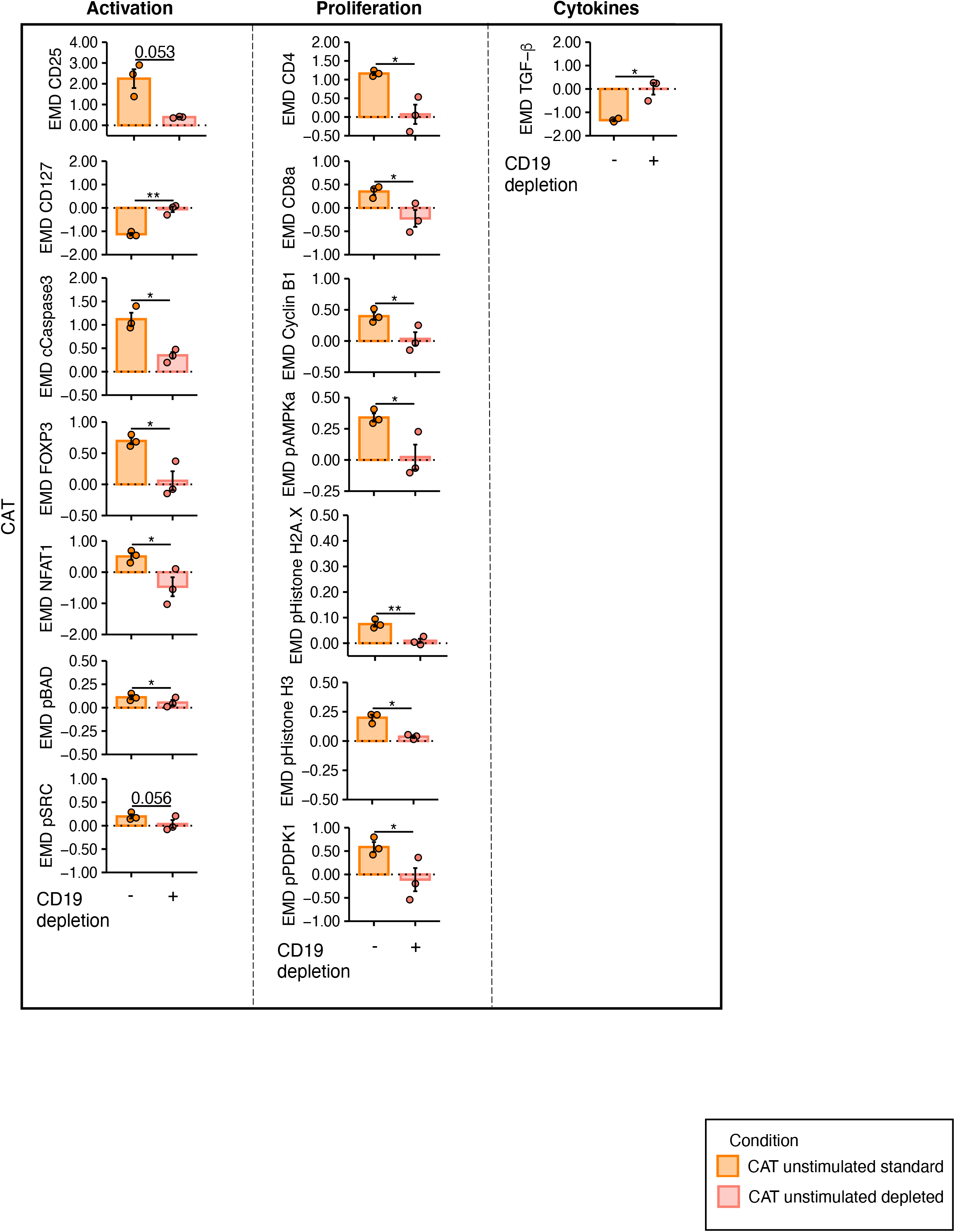
Mass cytometry EMD scores in standard or CD19-depleted unstimulated CAT CAR transduced T-cells. The barplots show mass cytometry EMD scores for activation/proliferation markers and cytokines differentially expressed between CAT CAR T-cells manufactured in standard or CD19-depleted conditions. The data shown are normalized to unstimulated CD3 UNTR T cell control. The dotted horizontal line (0) represents the expression of a specific marker in the unstimulated UNTR condition (n = 3 HDs, n = 1 independent experiment). Data represent mean ± se. Statistical significance was calculated by Paired t-test. *P < 0.05, **P < 0.01. Each experimental condition is indicated by a specific colour code (Standard condition orange; CD19-depleted condition salmon).

**Supplementary Fig. 9:**
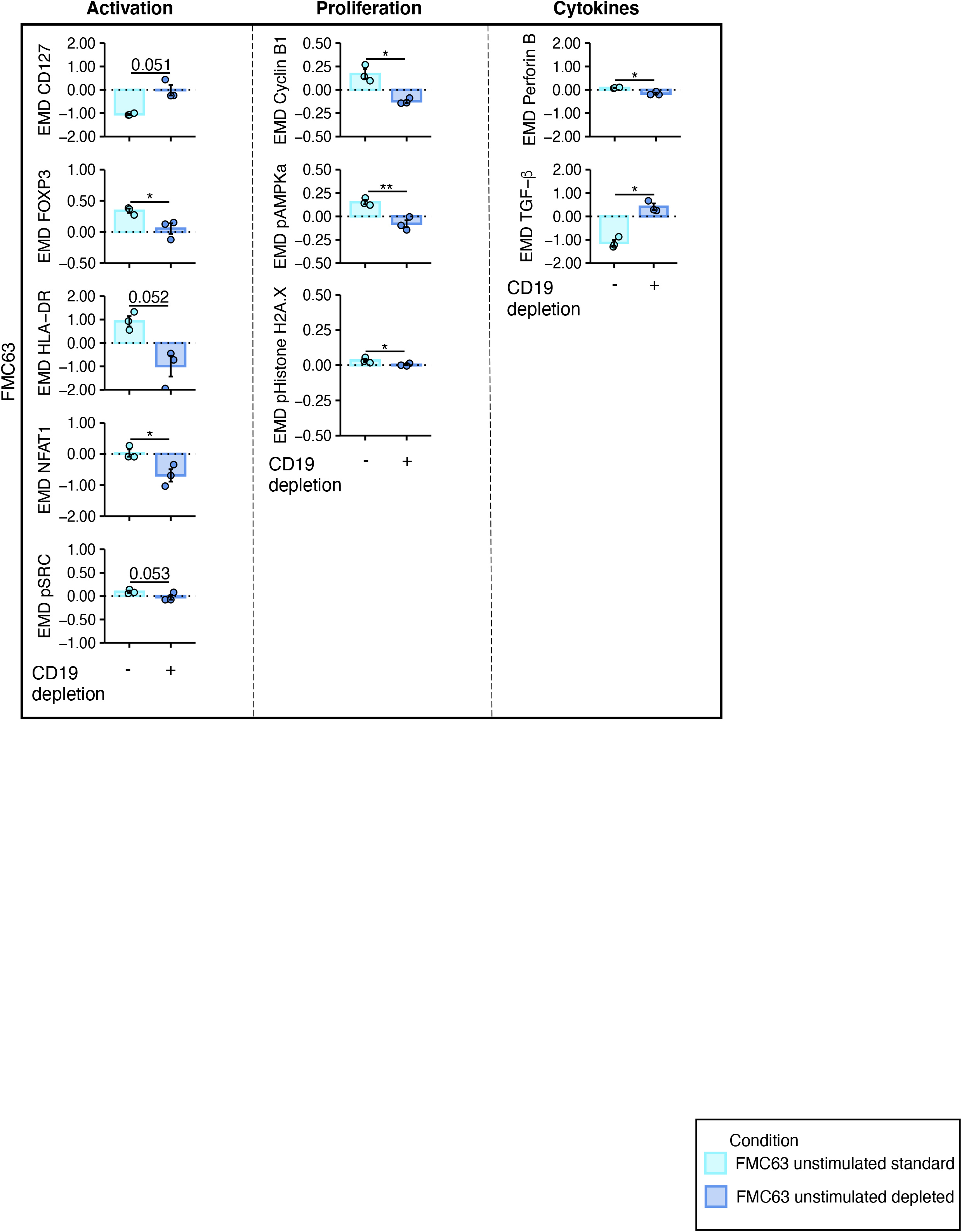
Mass cytometry EMD scores in standard or CD19-depleted unstimulated FMC63 CAR transduced T-cells. The barplots show mass cytometry EMD scores for activation/proliferation markers and cytokines differentially expressed between FMC63 CAR T-cells manufactured in standard or CD19-depleted conditions. The data shown are normalized to unstimulated CD3 UNTR T-cell control. The dotted horizontal line (0) represents the expression of a specific marker in the unstimulated UNTR condition. (n = 3 HDs, n = 1 independent experiment). Data represent mean ± se. Statistical significance was calculated by Paired t-test. *P < 0.05, **P < 0.01. Each experimental condition is indicated by a specific colour code (Standard condition light blue; CD19-depleted condition cornflower blue).

**Supplementary Fig. 10:**
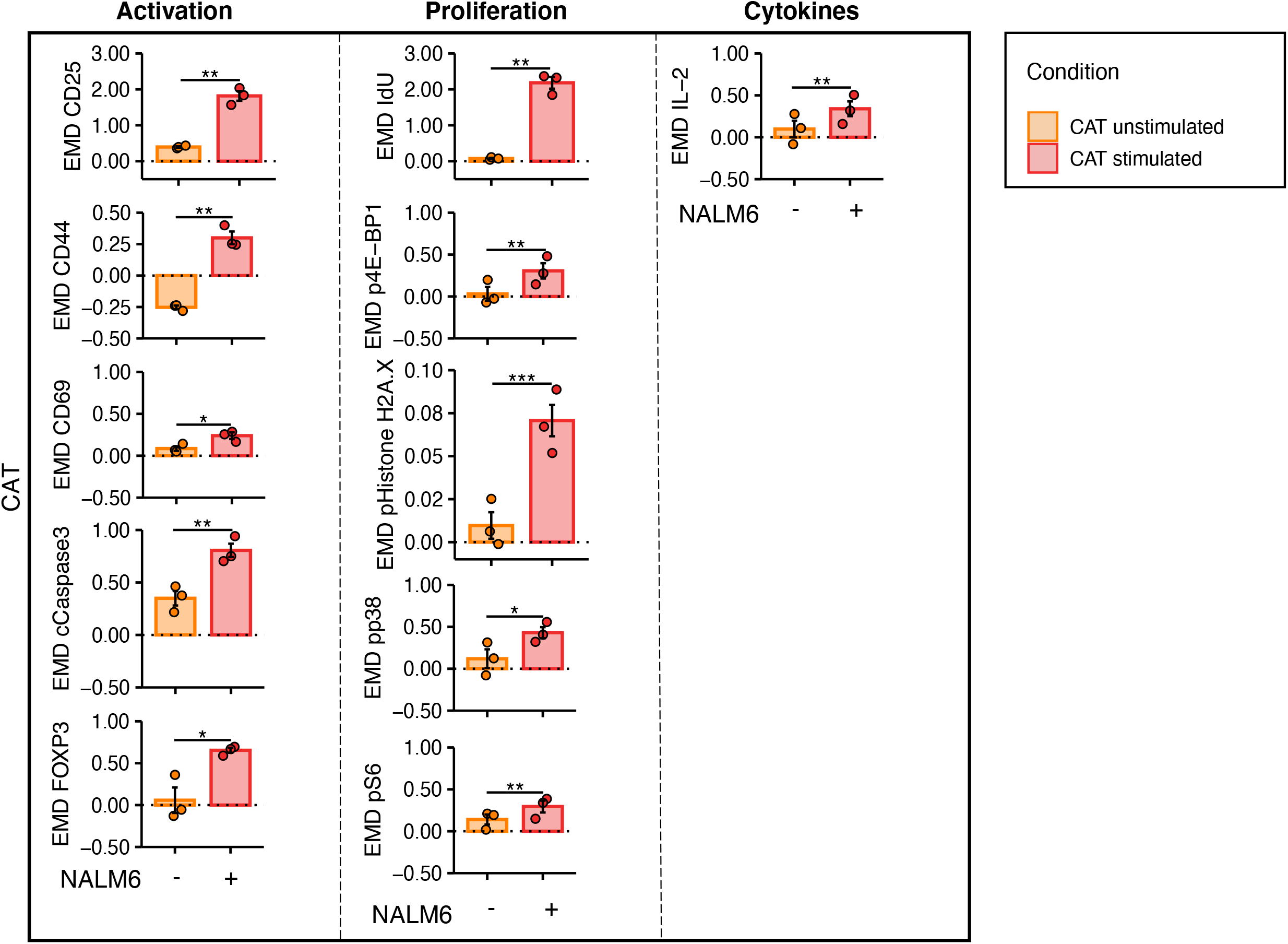
Mass cytometry EMD scores in unstimulated and stimulated CD19-depleted CAT CAR transduced T-cells. The barplots show mass cytometry EMD scores for activation/proliferation markers and cytokines differentially expressed between unstimulated and stimulated CAT CAR T-cells manufactured in CD19-depleted conditions. The data shown are normalized to unstimulated CD3 UNTR T-cell control. The dotted horizontal line (0) represents the expression of a specific marker in the CD19-depleted UNTR condition. (n = 3 HDs, n = 1 independent experiment). Data represent mean ± se. Statistical significance was calculated by Paired t-test. *P < 0.05, **P < 0.01, ***P < 0.001. Each experimental condition is indicated by a specific colour code (Unstimulated condition orange; stimulated condition red).

**Supplementary Fig. 11:**
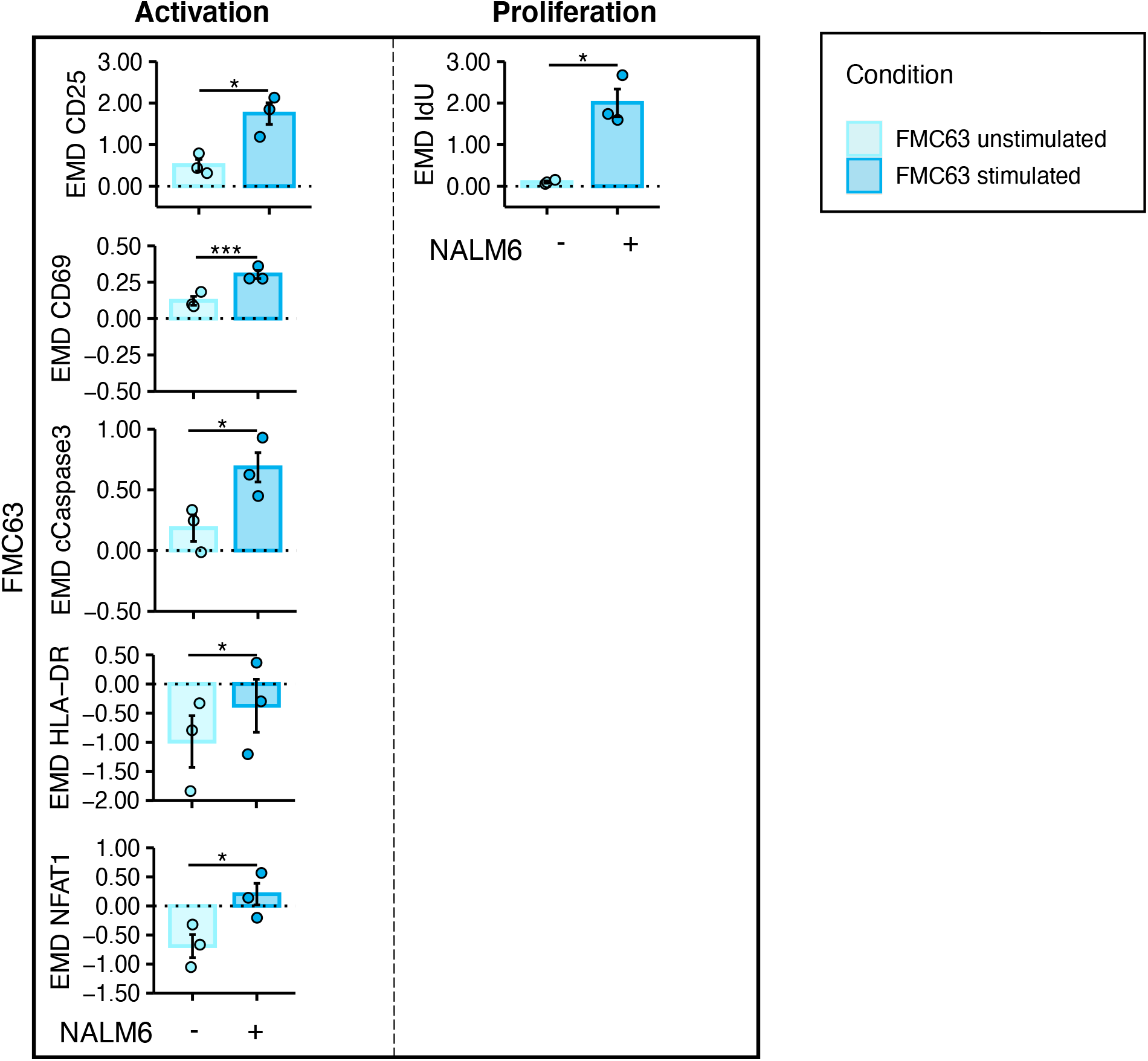
Mass cytometry EMD scores in unstimulated and stimulated CD19-depleted. **FMC63 CAR transduced T-cells.** The barplots show mass cytometry EMD scores for activation and proliferation markers differentially expressed between unstimulated and stimulated FMC63 CAR T-cells manufactured in CD19-depleted conditions. The data shown are normalized to unstimulated CD3 UNTR T-cell control. The dotted horizontal line (0) represents the expression of a specific marker in the UNTR CD19-depleted condition (n = 3 HDs, n = 1 independent experiment). Data represent mean ± se. Statistical significance was calculated by Paired t-test. *P < 0.05, ***P < 0.001. Each experimental condition is indicated by a specific colour code (Unstimulated condition light blue; stimulated condition blue).

## Acknowledgements

We are grateful to the UCL ICH Flow Cytometry facility for support in cell-sorting. We are grateful to Dr Thomas Adejumo (Fluidigm) for mass cytometry valuable suggestions and assistance with protocols design. We thank Dr Anne Marijn Kramer (Amsterdam UMC) for generating CD19 FMC63 and CAT CAR transfer vector plasmids with S.G. and M.P. and P.J.A. The authors acknowledge the contribution of UCL Genomics Facility. This work was supported by the NIHR GOSH BRC (NIHR GOSH BRC 17PA01), the views expressed are those of the author(s) and not necessarily those of the NHS, the NIHR, or the Department of Health. Part of this work was supported by the Leukaemia UK John Goldman Fellowship to A.G. (2018/JGF/003), the Rosetrees Trust fund to A.G. (M700) and the Academy of Medical Sciences Springboard Award to A.G. (SBF004\1025).

## Author contributions

I.M.M. designed, performed and analyzed experiments, performed bioinformatic analyses of mass cytometry experiments and contributed to writing the manuscript. E.G-C performed bioinformatic analyses of transcriptomic data and wrote the relative bioinformatic supplementary information. R.V.C.P. performed experiments and bioinformatic analyses of mass cytometry and transcriptomic data. F.C-R provided data analysis tools and contributed to bioinformatic analyses of mass cytometry experiments. P.P-C. performed transcriptomic bioinformatic analyses data checks and contributed to writing the bioinformatic supplementary information. J. S., M.S., S.W.W, A.Gu. and E.K. performed experiments. A.E. provided support to cell sorting. J.F. provided analytical pipelines and useful discussion for the analysis and normalization of mass cytometry data. S.G. and M.P. provided CAR constructs. C.J.T provided expertise in mass cytometry and reagents. P.J.A. provided reagents and expertise and contributed to writing the manuscript. S.C. supervised the bioinformatic analyses and contributed to writing the manuscript. A.G. designed and supervised the project, performed and analyzed experiments and wrote the manuscript. All authors provided critical feedback and helped shape the research, analysis and manuscript.

## Correspondence

Corrispondence to Dr Alice Giustacchini.

